# Molecular basis for inhibition of adhesin-mediated bacterial-host interactions through a novel peptide-binding domain

**DOI:** 10.1101/2020.11.18.389114

**Authors:** Shuaiqi Guo, Hossein Zahiri, Corey Stevens, Daniel C. Spaanderman, Lech-Gustav Milroy, Christian Ottmann, Luc Brunsveld, Ilja K. Voets, Peter L. Davies

## Abstract

Modulation of protein-protein interactions (PPIs) with small-molecules is a promising conceptual approach in drug discovery. In the area of bacterial colonization, PPIs contribute to adhesin-mediated biofilm formation that cause most infections. However, the molecular basis underlying these adhesin-ligand interactions is largely unknown. The 1.5-MDa adhesion protein, *MpIBP,* uses a peptide-binding domain (*Mp*PBD) to help its Antarctic bacterium form symbiotic biofilms on sea ice with microalgae such as diatoms. X-ray crystallography revealed *Mp*PBD uses Camdependent interactions to self-associate with a crystal symmetry mate via the C-terminal threonine-proline-aspartate sequence. Structure-guided optimization derived penta-peptide ligands that bound *Mp*PBD 1,000-fold more tightly, with affinities in the nano-molar range. These ligands act as potent antagonists to block *Mp*PBD from binding to the diatom cells. Since adhesins of some human pathogens contain peptide-binding module homologs of *Mp*PBD, this same conceptual approach could help develop ligand-based PPI modulators to disrupt harmful bacteria-host interactions.

## Introduction

Protein-protein interactions (PPIs) are central to most biochemical processes such as actin and tubulin polymerization, ATP production through the electron transport chain, and signal transduction via G protein-coupled receptors(1–3). Given PPIs’ vital role in the well-being of cells, their modulation by drug-like molecules is an attractive approach that holds great promise for the development of new strategies to treat various diseases. Over the past two decades, significant advances in structural biology and medicinal chemistry have enabled some fruitful developments in drug discovery to target PPIs that govern mammalian cellular processes(4–6). For example, the structure of B-cell lymphoma extra-large (Bcl-xL) protein in complex with a peptide derived from the Bcl-2 Antagonist/Killer (Bak) protein inspired the fragment-based identification of the first Bcl-2 family inhibitors(7). This ultimately led to the FDA approval of Venetoclax in 2016 to treat lymphocytic leukemia and lymphoma(8). In addition to such small-molecule inhibitors, peptide antagonists have been developed to target their partner proteins’ ligand-binding sites to disrupt PPIs of Bcl-2 proteins(9) and those involved in other diseases(10–13). Although the use of small-molecule and peptide therapeutics to target PPIs is an active and highly promising field(12–14), most of the studies published to date have been focused on cancer-related, human PPIs. Investigations of PPIs that mediate bacterial infections have taken second place to antibiotic development. It is only now when antibiotics are losing some of their potency that other strategies for inhibiting bacteria are being intensively studied(15–18).

Bacterial adhesins are a key class of virulence factors that bind bacteria to host cells(19–22), and subsequently help develop multi-cellular communities called biofilms responsible for over 80% of chronic infections in humans(23). There is currently a shortage of effective treatments against biofilm-related infections(23–25). With the increasing spread of antibiotic-resistant pathogenic bacteria, there is an urgent need for the development of PPI disruptors to block the adhesins of bacteria, which might prevent the colonization and persistent infections caused by biofilms. Adhesins are typically long, modular proteins with one terminus anchored to the bacterial surface, while the other end is extended out to interact with substrates like the carbohydrates and proteins on host cell surfaces(19, 26–31). With structures of adhesin-ligand complexes starting to emerge(32–34), researchers are beginning to elucidate the molecular basis of the interactions required to build a biofilm. To date, most of these studies have focused on the characterization of lectin-glycan interactions(32, 33, 35), and have resulted in some effective treatments against bacterial infections(17, 36). For example, the FimH adhesin of uropathogenic *Escherichia coli* has been successfully targeted by mannose analogs to help treat urinary tract infections(35, 37–41). These studies validate the efficacy of the anti-adhesion strategy, which can be used as an alternative approach to treat bacterial infections without the excessive use of antibiotics. While PPIs are heavily involved in bacteria-host interactions, examples that detail the molecular basis of these interactions are scarce. Given that PPIs are typically many fold stronger than those of lectin-glycan interactions(42), the development of PPI inhibitors to disrupt bacteria-host interactions will be of great interest and potential utility.

*Marinomonas primoryensis* ice-binding protein (*Mp*IBP) is an exceptionally large (~ 1.5 MDa) Repeats-In-Toxin (RTX) adhesin found on the surface of its Antarctic bacterium(27, 43). While its N-terminal Region I (RI) is responsible for anchoring the adhesin to the outer-membrane, its C-terminal ligand-binding Regions III and IV (RIII and RIV) help the bacterium form symbiotic biofilms with diatoms on the underside of lake ice(27, 28). *Mp*IBP_RIII contains five β-sheet-rich domains, including a carbohydrate-binding lectin module (RIII_5, also referred as *Mp*PA14) and a Peptide-Binding Domain (RIII_3, *Mp*PBD), both responsible for binding diatoms by interacting with the cell-surface glycans and proteins, respectively. We reported the X-ray crystal structure of *Mp*IBP_RIII1-4, which revealed that *Mp*PBD folds as an oblong β-sandwich with a shallow, solvent-exposed ligand-binding cavity on the periphery of the structure(27). Importantly, it was observed in the crystal unit cell that the three C-terminal amino-acid residues (threonine-proline-aspartate, TPD) of *Mp*IBP_RIII1-4 were stably anchored in the ligand-binding cavity of a symmetry-related molecule within the crystal unit cell. The observed crystal contacts led us to postulate that *Mp*PBD binds its bacterium to diatoms by interacting with surface proteins via short tripeptide sequences at the C-termini.

Here, we show that the TPD peptide sequence binds *Mp*PBD in solution with an affinity of 26 μM. Using a structure-guided approach, we optimized the peptidyl sequence by iteratively screening a library of pentapeptides and obtained ligands that bound *Mp*PBD ~ 1000-fold more strongly with affinities of ~ 30 nM. X-ray crystal structures of *Mp*PBD-ligand complexes revealed that the strong protein-peptide interactions originate from a combination of Ca^2+^-dependent polar interactions and hydrophobic contacts. We further demonstrate that these short peptidyl ligands are effective at inhibiting the binding of *Mp*PBD to diatoms, which gives insight into how microbial adhesion can be disrupted through ligand-based antagonists.

## Results and discussion

### Initial ligand identification

Close inspection of the *Mp*IBP_RIII1-4 structure (PDB code: 5K8G) revealed the detailed PPIs involved in its self-association within the crystal by binding the C-terminal Thr-Pro-Asp (TPD) of a symmetry-related molecule (Figure 1A). The C-terminal aspartate residue is buried inside the ligand-binding pocket of the other molecule, with its terminal α-carboxyl oxygens binding Ca1 and Ca2 via three ionic bonds (Figure 1B). The protein-protein interaction is further enhanced by hydrogen bonding between carbonyl and amide groups of the TPD sequence and V293 and N333 from the Calcium- and Peptide-Binding Loops 1 and 2 (CPBL1 and 2) of *Mp*PBD. The threonine residue at the pre-penultimate position (position 3) is the most solvent exposed residue of the peptide, and the ambiguous electron density of its side chain was indicative of a high degree of flexibility. The fortuitous crystallisation of the self-interacting *Mp*PBD_RIII1-4 indicated that peptidyl sequences are the likely ligands of *Mp*PBD. To prevent the self-association through the extended C-terminal “TPD” sequence, and facilitate the binding of free peptide to *Mp*PBD, we designed a new *Mp*IBP_RIII1-4 construct by truncating the carboxyl end of the original protein from D519 to N507 (Figure 1A, Supplementary file 1-Figure S1; green and blue arrows). This new *Mp*IBP_RIII1-4 construct was used for the subsequent binding and structural studies described below.

**Figure 1:**
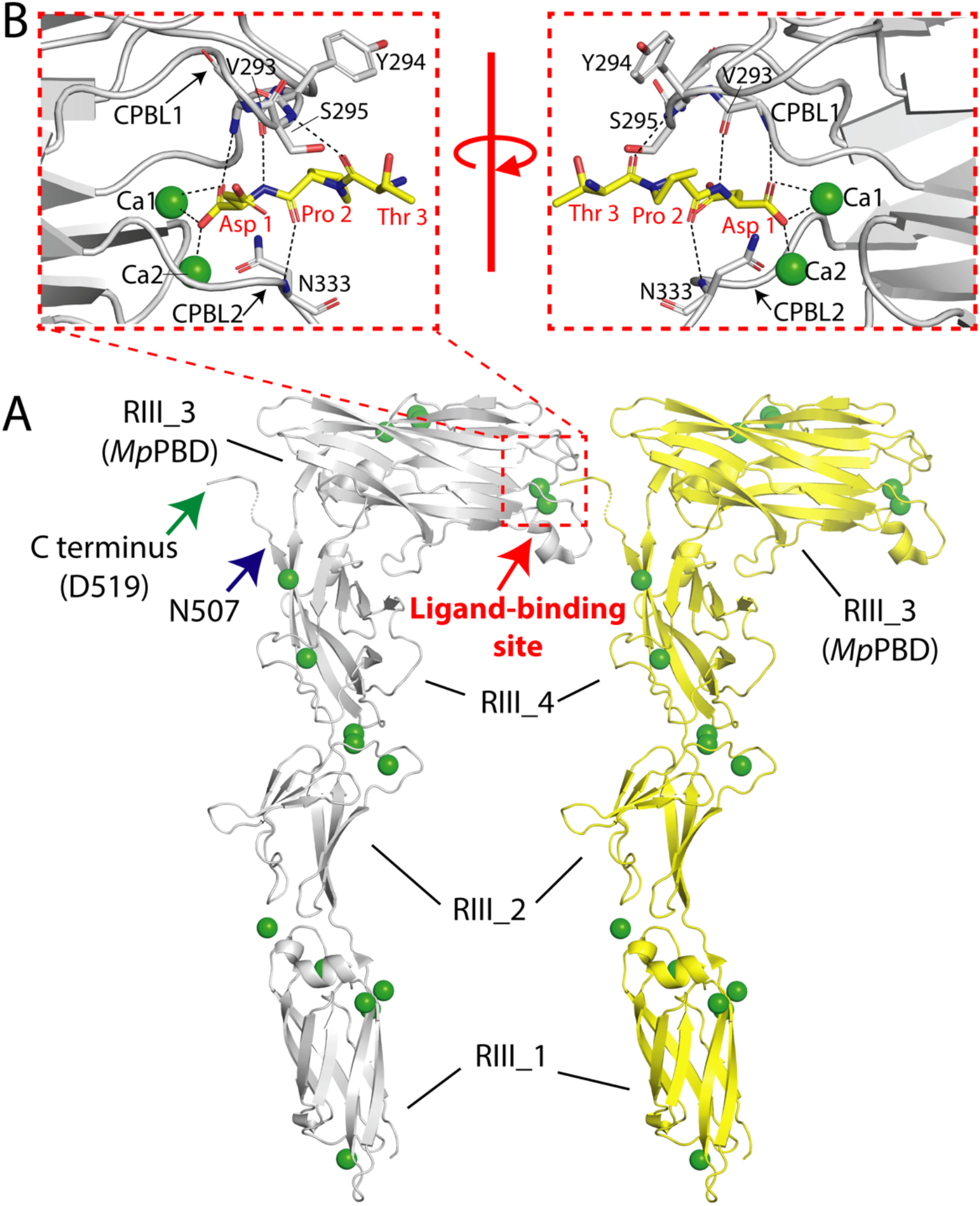
Self-association of *Mp*IBP_RIII1-4 within the crystal unit cell. A) *Mp*IBP_RIII1-4 molecule (grey) interacts with a neighboring symmetry mate (yellow) in the crystal via its C-terminal “TPD” sequence. The C-terminal D519 residue of the original *Mp*IBP_RIII1-4 construct is marked by a green arrow, while the C-terminal N507 residue of the truncated construct used in this study is marked by a blue arrow. The ligand-binding site is indicated by a red box of dashed lines. B) Zoomed-in view of the ligand-binding site of *Mp*RIII1-4 within the crystal showing the atomic details at the protein-protein interaction interface. The right panel shows the interface from a view that is rotated approximately 180° around a vertical axis from the left. Polar interactions are indicated by black dashed lines. Carbon atoms of the bound “TPD” sequence are colored in yellow while those for its symmetry-related protein are colored in grey. Oxygen atoms are red, nitrogen atoms are blue and Ca^2+^ ions are shown as green spheres. Amino acid residues involved in protein-protein interactions are labelled and shown in stick representation.

Given that a C-terminal carboxylic acid group appears to be key for the proteinprotein interaction, we used fluorescence polarisation (FP) to screen a small collection of 15 N-terminally FITC-labelled peptides that end in various C-terminal amino acid residues with a free α-carboxyl group (selected on the basis of availability within the laboratory). In the presence of ~ 80 μM of *Mp*IBP_RIII1-4, two FITC-labelled and phosphorylated peptides with sequences of IKARAS(pS)SPVILVGTHLD (pep 14) and RHKKLMFK(pT)EGPDSD (pep 15) produced significant delta fluorescence polarisation values, indicating binding of these peptides to *Mp*PBD (Figure 2A). None of the other 13 peptides appeared to bind *Mp*PBD as they produced negligible delta polarisation values. FP with protein titrations further showed that in the presence of 2 mM Ca^2+^, peptides 14 and 15 bound *Mp*PBD with EC_50_ values of 2.7 μM and 0.69 μM, respectively. Moreover, the presence of excess EDTA abolished the interaction between peptide and protein (Figure 2B), which validated the Ca^2+^-dependency of the protein-peptide interaction as demonstrated by the structural data.

**Figure 2:**
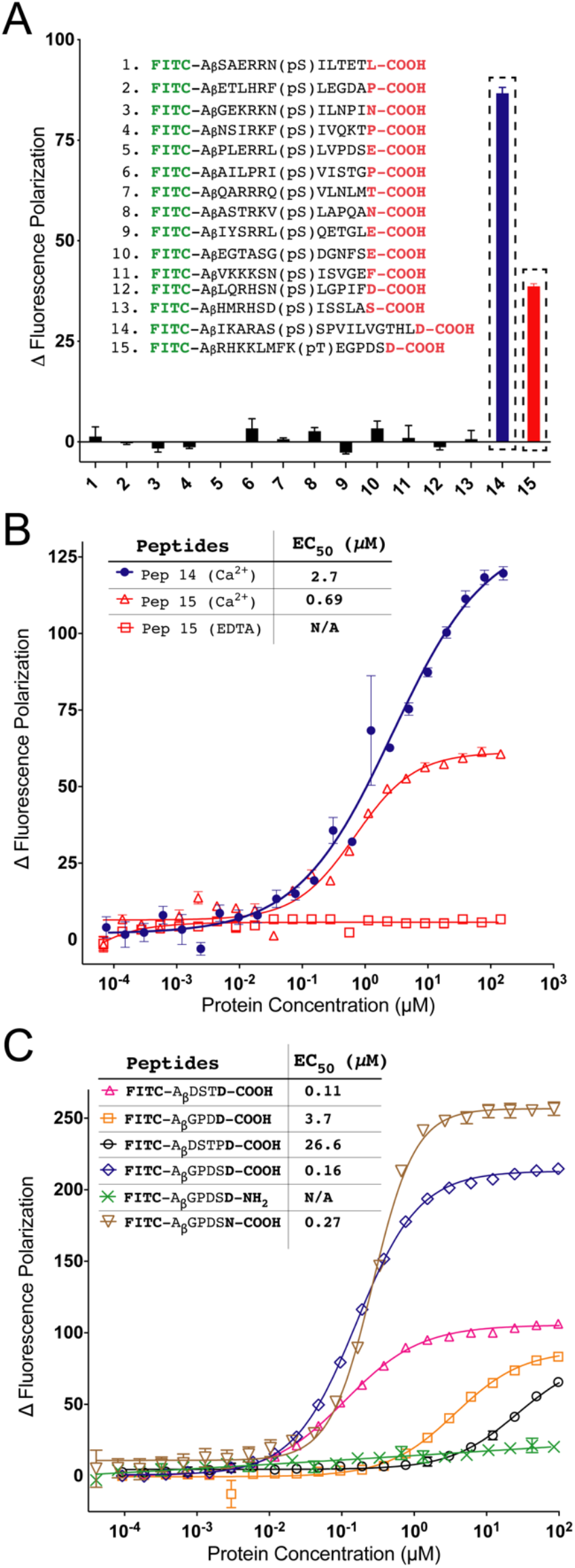
Identification of initial peptidyl ligands of *Mp*PBD. (A) Screening of a collection of 15 FITC-labelled peptides by fluorescence polarisation (FP) with *Mp*IBP_RIII1-4 at a concentration of ~ 80 μM. Background polarization was subtracted from all values to result in Δ polarisation values on the y-axis. Peptide sequences are displayed. Data bars for peptides 14 (blue) and 15 (red) are indicated within boxes of dashed lines. (B) FP assay of Peptides 14 (blue closed circles) and 15 (red open triangles) from (A) with titration of *Mp*IBP_RIII1-4 in the presence of 2 mM CaCl_2_ and in the presence of excess 2 mM EDTA (red open squares). (C) FP assay to assess the binding of six different FITC-labelled peptides to *Mp*PBD. The mean of three experiments were plotted. Some of the SD error bars are smaller than the data point symbols in B) and C). The calculated EC_50_ values for each experiment are shown in B) and C).

Interestingly, the three binding sequences “TPD”, pep 14 and pep 15 all ended in a C-terminal aspartate residue with a free α-carboxylic acid group. This was consistent with the structural data, which showed that the terminal carboxylic acid group of “TPD” directly bonded to the two calcium ions in the *Mp*PBD ligand-binding site. However, since the C-terminal aspartate side chain of *Mp*IBP_RIII1-4 had no direct interactions with its neighbouring symmetry mate in the crystal, it was not clear how the aspartate residues in these three peptidyl sequences contributed to their binding to *Mp*PBD. To further validate the hypothesis that *Mp*PBD binds specifically to certain C-terminal amino-acid sequences, peptides encompassing only the last five amino acids of the original *Mp*IBP_RIII1-4 construct (FITC-A_β_DSTPD) and pep 15 (FITC-A_β_GPDSD) were synthesized and studied. The affinity of *Mp*PBD for these two short peptide ligands, together with two analogs (FITC-A_β_DSTD and FITC-A_β_GPDD) was measured by FP (Figure 2C).

Unexpectedly, FITC-A_β_DSTPD, which contains the C-terminal “TPD” binding sequence originally identified in the crystal structure of *Mp*IBP_RIII1-4, showed the weakest interaction to *Mp*PBD out of this series, with an EC_50_ of ~ 27 μM. In contrast, the peptide, FITC-A_β_DSTD, bound at least 200-fold stronger with an EC_50_ of 0.11 μM (Figure 2C). Similarly, FITC-A_β_GPDSD bound with an EC_50_ of 0.16 μM, which is roughly 4-fold stronger than its longer, phosphorylated counterpart (FITC-RHKKLMFKpTEGPDSD, EC_50_ = 0.69 μM), validating the importance of the C-terminal residues for binding *Mp*PBD. Furthermore, FITC-A_β_GPDSD bound the protein 20-fold stronger than its analog FITC-A_β_GPDD.

To further ascertain the chemical constituents of the peptide C-terminus responsible for binding *Mp*PBD, two additional analogs of the peptide FITC-A_β_GPDSD were synthesized. When the C-terminal aspartate was replaced by an asparagine, a residue with similar length side chain but different chemistry, the affinity of the peptide FITC-A_β_GPDSN for *Mp*PBD fell by 40% to an EC_50_ of 0.27 μM (Figure 2C). However, the substitution of the terminal α-carboxylic acid group of FITC-A_β_GPDSD by a C-terminal carboxamide group (FITC-A_β_GPDSD-NH_2_) abolished the protein-peptide interaction all together (Figure 2C). These results demonstrated the crucial importance of the ionic interaction between the terminal carboxylic acid group and the two Ca^2+^ ions in the ligand-binding site of *Mp*PBD. The side-chain carboxylic acid group of the peptide C-terminal aspartic acid residue gave more favourable binding to *Mp*PBD compared to that of the side-chain amide of the peptide bearing an asparagine at the same position. Moreover, the FP results indicate that the peptide amino acids at the penultimate position (position 2) have a significant impact on the peptide-protein interactions. The strong binders, FITC-A_β_GPDSD and FITC-A_β_DSTD, contain the small polar residues of either threonine or serine at this position. The intermediate binder FITC-A_β_GPDSD has an aspartate while the weakest binder FITC-A_β_DSTPD has a proline at the same position. Although the proline is involved in main-chain hydrogen bonding with V293 and N333 of the CPBL1 and CPBL2, respectively, its rigid five-membered imine ring is unable to interact with the protein (Figure 1B). As for FITC-A_β_GPDD, the negative charge of the aspartate side chain at position 2 may be disruptive for the peptide-protein interaction. We therefore sought to acquire detailed structural information to explain the high affinities of FITC-AβGPDSD and FITC-A_β_DSTD for *Mp*PBD.

### Structural basis for peptide-MpPBD interactions

To elucidate the atomic details of the peptide-protein interactions, we solved the X-ray crystal structures of *Mp*IBP_RIII1-4 in complex with the peptides FITC-A_β_DSTD and FITC-A_β_GPDSD to 2-Å resolution (Supplementary file 1-Table S1). Clear electron densities were observed for the peptide amino acids in the last three positions (Figure 3), while those at the position immediately beforehand at the N-terminal end appeared to be ambiguous. No electron density could be observed beyond position 3 suggesting these residues are highly flexible because they are fully exposed to the solvent and are thus unlikely to interact with the protein.

**Figure 3:**
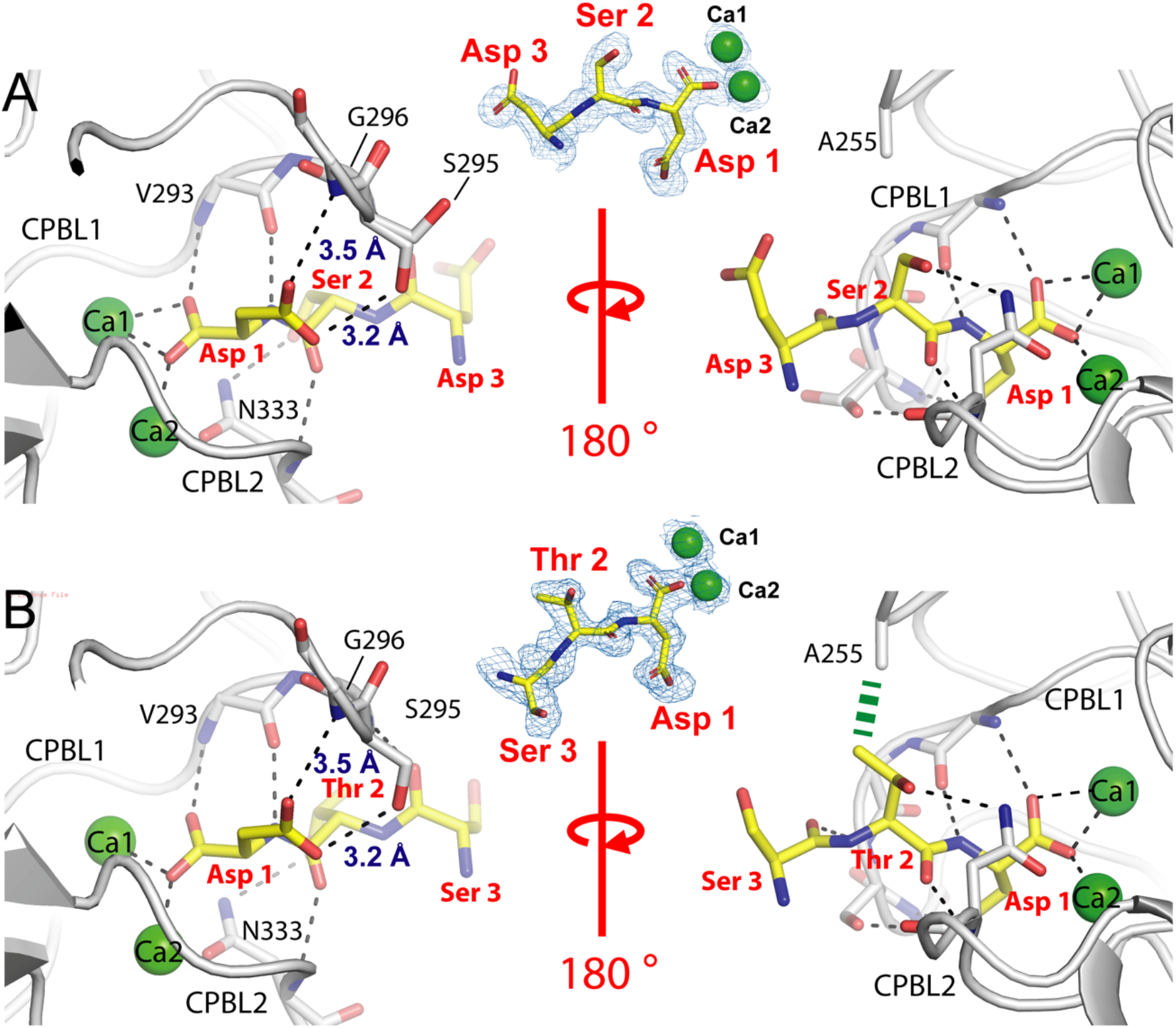
Protein-peptide binding interfaces revealed by X-ray crystallography. Zoomed-in views of the *MpPBD* ligand-binding site interacting with peptides that end with sequences of DSD (A) and DSTD (B), respectively. The 2Fo – Fc electron density maps around the peptide residues and Ca1 and 2 are shown as blue mesh (contoured at 1 σ). The right panels in A) and B) are views that are rotated around approximately 180° from the left panels. Polar interactions are indicated by black dashed lines. Carbon atoms of the peptides are colored in yellow while those for the protein are colored in grey. Oxygen atoms are red, nitrogen atoms are blue, while Ca^2+^ ions are shown as green spheres. Amino acids involved in protein-peptide interactions are shown in stick representation. Peptide amino-acid residues are indicated in red 3-letter code, while those for the protein are labelled in black one-letter code. CIF files can be found in Figure 3 – source data 1 and has been deposited in the Protein Data Bank (PDB: “Code”)

At position 2 of FITC-A_β_GPDSD, serine has its side-chain hydroxyl group hydrogen bonded with the side-chain amide of N333 on CPBL2, while the main chains of these two residues hydrogen bond via their carbonyl and amide groups (Figure 3A). The observed hydrogen bonding interactions help to explain why FITC-A_β_GPDSD has a 20-fold higher affinity for *Mp*PBD than that of FITC-A_β_GPDD, with the only difference between the two peptides being the presence or absence of serine at position 2. Indeed, X-ray crystallography indicates that the larger aspartate residue at position 2 would clash with N333 and destabilize the protein-peptide interactions. In comparison, FITC-A_β_DSTD has a similar binding mode to FITC-A_β_GPDSD, with the threonine side-chain hydroxyl hydrogen bonded with the amide of N333 side chain at the edge of the peptide-binding cavity. In addition, the threonine side-chain methyl group is involved in hydrophobic contact with residue A255 of the protein (Figure 3B). This helps restrain the free rotation of the threonine side-chain hydroxyl, locking it into a favourable conformation for polar interactions. The observed additional hydrophobic interactions might help explain why FITC-A_β_DSTD bound *Mp*PBD with slightly higher affinity than did FITC-A_β_GPDSD.

Interestingly, the gain of hydrogen bonding interactions at the peptide 2^nd^ position had an impact on how the C-terminal aspartate of the peptides bound *Mp*PBD. The aspartate of the TPD sequence observed in the ligand-binding cavity showed no interaction between its side chain γ carboxylic acid group and the protein. In contrast, when FITC-A_β_DSTD and FITC-A_β_GPDSD were complexed with *Mp*PBD, the side-chain hydroxyl of S295 within CPBL1 pointed downward to hydrogen bond with the γ carbonyl of the peptide aspartate, which helped G296 to interact with the γ hydroxyl of the same peptide aspartate via its main-chain amide (Figure 3A, B, left panels). Taken together with the gained interactions from the peptide position 2, the structural data gave the molecular explanation of why FITC-A_β_DSTD and FITC-AβGPDSD bound *Mp*PBD with substantially higher affinity than their analogs, FITC-AβDSTPD and FITCAβGPDD, respectively.

### Structure-guided screening identified ligands with low-nano-molar affinity

Guided by the structural insight that the C-terminal residues are crucial for the peptide-protein interactions, we next sequentially screened libraries of pentapeptides by FP to identify stronger ligands for *Mp*PBD.

The first round of screening involved 20 pentapeptides with a consensus sequence of FITC-AGAGX where the first four amino acids of the peptides were alternating alanine and glycine residues. The ultimate (C-terminal) residue “X” represented one of the 20 naturally occurring amino acids (Figure 4A). Consistent with the results shown above, the peptide with an aspartate at the C-terminal position (FITC-AGAGD) bound *Mp*PBD the tightest, with a moderate EC_50_ of 3.2 μM. This was followed by those peptides with large hydrophobic side-chains at position 1, such as FITC-AGAGI, FITC-AGAGY and FITC-AGAGF, which produced EC_50_ values of 4.4 μM, 4.6 μM, and 7.2 μM, respectively, when bound to *Mp*PBD (Supplementary file 1-Table S2, Figure S2). Most of the other peptides bound *Mp*PBD with an affinity in the 10 μM range, while those with basic side chains (FITC-AGAGK and FITC-AGAGR) had negligible interaction with *Mp*PBD.

**Figure 4:**
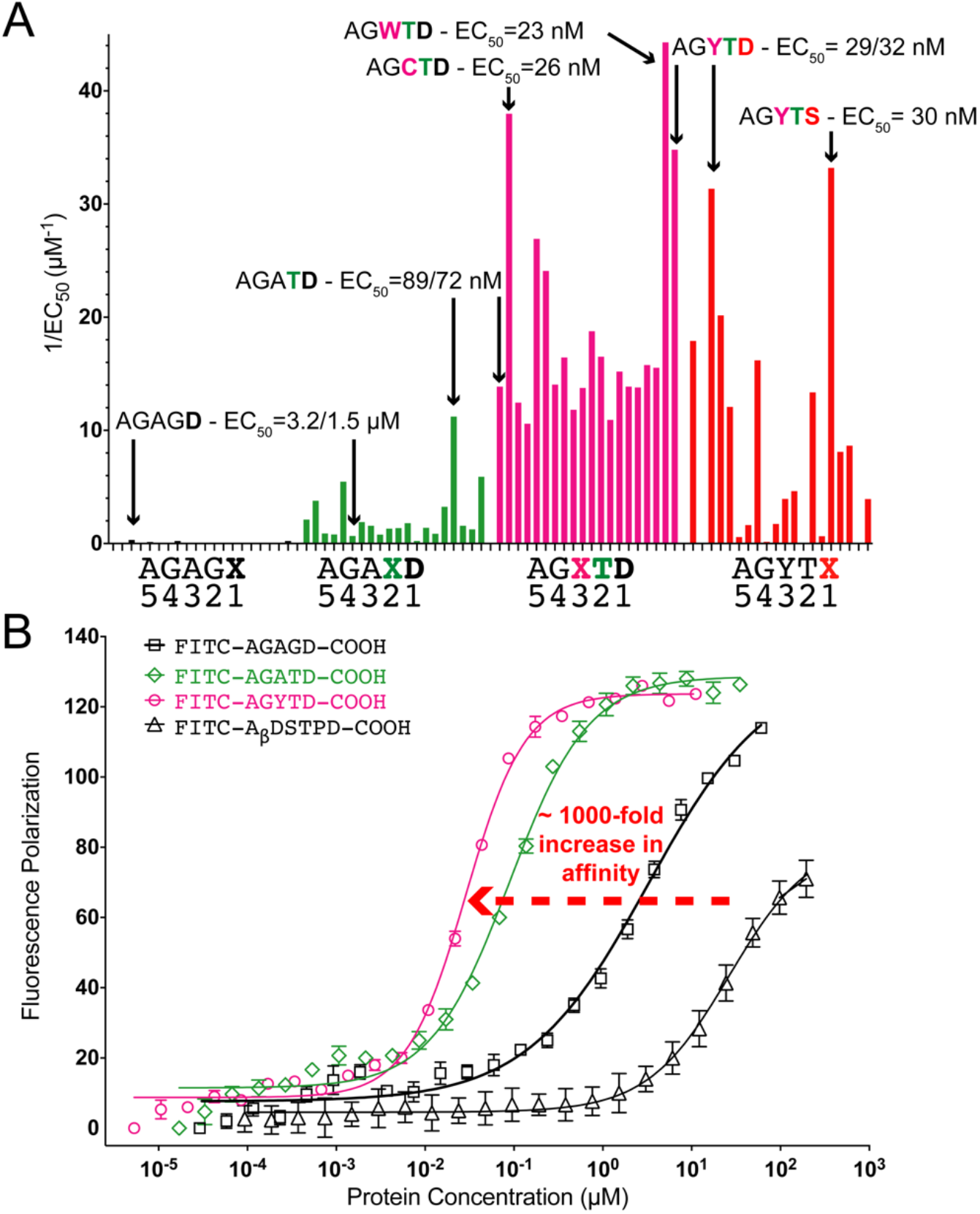
Structure-guided optimization of peptidyl ligands of *MpPBD.* (A) Overview of the binding of the 80 FITC-labelled pentapeptides to *Mp*PBD in four rounds of screening. Data are plotted as average 1/EC_50_ values calculated from FP assays of each individual FITC-labelled peptide (in three replicates). All EC_50_ values are listed in Tables S2, S3, and the corresponding FP titration plots are shown in Figures S2-S5. Data bars for peptides in the first round of screening (FITC-AGAGX) are colored black, while those for second (FITC-AGAXD), third (FITC-AGXTD) and fourth (FITC-AGYTX) rounds are colored green, magenta and red, respectively (only some amino-acid sequences are labelled in the graph for short). The strongest *Mp*PBD binders from each round of screening are marked by arrows and their calculated EC_50_ values are indicated. (B) FP titration plots of representative peptides showing the progressive enhancement of ligands’ affinity for *Mp*PBD.

Having identified FITC-AGAGD as the strongest ligand for *Mp*PBD in the first round of screening, we proceeded with a second set of 20 pentapeptides that had a consensus sequence of FITC-AGAXD. All peptides except for AGAPD (EC_50_ = 4.6 μM) bound more strongly than FITC-AGAGD (EC_50_ =1.5 μM). Consistent with the structural studies reported earlier (Figure 1B), neither proline nor glycine at position 2 can have side-chain interactions with *Mp*PBD, explaining their lower affinity for the protein. Furthermore, the peptide with threonine in the second position (FITC-AGATD) has the highest affinity for *Mp*PBD with an EC_50_ of 89 nM (Figure 4A, green; Supplementary file 1-Table S2; Figure S3), which is up to 35-fold tighter than those of FITC-AGAGD. Peptides with aromatic residues at the same position, including FITC-AGAFD and FITC-AGAYD bound slightly weaker than did AGATD, with EC_50_ values of 180 nM and 170 nM, respectively. Consistent with the characterisation of FITC-A_β_GPDSD and FITC-A_β_DSTD, FITC-AGATD bound *Mp*PBD more than 3-fold stronger than did FITC-AGASD (EC_50_ = 310 nM), validating the importance of the threonine methyl group in restraining the hydroxyl group to a favourable conformation for interacting with N333.

With the last two C-terminal residues defined as Thr and Asp, we proceeded to screen for the optimal amino acid in the 3^rd^ position from the C terminus (FITC-AGXTD). While the majority of the 20 peptides of FITC-AGXTD bound *Mp*PBD more strongly than did FITC-AGATD (72 nM in this round of screening), with affinity in the nanomolar range, the three with aromatic side chains and cysteine stood out. In particular, FITC-AGYTD, FITC-AGWTD and FITC-AGCTD produced EC_50_ values of 29 nM, 23 nM, and 23 nM (Figure 4A, magenta; Supplementary file 1-Table S3; Figure S4), respectively. We reason that these amino acids with hydrophobic side chains at the 3^rd^ position likely help the solvent-exposed residue interact with the protein via hydrophobic interactions, contributing to their higher affinity for *Mp*PBD.

Having demonstrated that residues at the 2^nd^ and 3^rd^ positions of the peptides have an impact on how the C-terminal amino-acid interacted with *Mp*PBD, we performed a final round of screening, with a consensus sequence of FITC-AGYTX (Figure 4A, red). The reasons for selecting the tyrosine-containing peptide sequence over the tryptophan-or the cysteine-containing peptide include: FITC-AGYTD is more soluble than FITC-AGWTD; FITC-AGYTD is not subject to dimerization the way FITC-AGCTD can be linked by cysteine-dependent disulphide formation, which might confound the results of the binding studies. The results of the screening revealed that all but one of the FITC-AGYTX peptides bound *Mp*PBD weaker than did FITC-AGYTD, with their EC_50_ values ranging from high nano-to micro-molar concentrations. This validated the importance of the C-terminal aspartate residue for the protein-peptide interaction. The only peptide with a comparable affinity to FITC-AGYTD is FITC-AGYTS, which produced a calculated EC_50_ of 30 nM (Figure 4A, red; Supplementary file 1-Table S3, Figure S5). Given their similar affinities for *Mp*PBD, AGYTD and AGYTS were considered two of the optimal peptidyl ligands, which bound *Mp*PBD roughly 1,000-fold tighter than the “TPD” binding sequence initially identified (Figure 4B).

### Isothermal titration calorimetry validated the nano-molar affinity of MpPBD ligands

To further validate the peptidyl ligands identified by FP, we used isothermal titration calorimetry (ITC) to directly measure the interactions between unlabelled peptides (i.e. lacking FITC labels) and *Mp*PBD. ITC measurements of *Mp*IBP_RIII1-4 (~ 20 μM) titrated with the four different ligands (~ 200 μM) of AGAGD, AGATD, AGYTD, and AGYTS all yielded sigmoidal-shaped curves with calculated stoichiometry values (N) close to 1 (Figure 5A-D). This was consistent with the X-ray crystallography data that demonstrated *Mp*PBD has only one ligand-binding site. The interaction of the moderate binder, AGAGD, with *Mp*PBD showed a gradually transitioning sigmoidal curve, indicating a relatively slow saturation rate of the *Mp*PBD ligand-binding sites. This resulted in a calculated K_d_ of 1.9 μM, which is comparable to the EC_50_ values of FITC-AGAGD obtained from the FP experiments (1.5 μM or 3.2 μM). The result suggested that the FITC label did not have a significant impact on the peptide-protein interactions. The three strong binders, AGATD, ATYTD and ATYTS, all resulted in sigmoidal-shaped ITC curves with much steeper transitions. The three peptides had calculated K_d_ values of 86 nM, 67 nM and 59 nM, respectively. Thus, the ITC results of the representative peptides showed the same trend in binding *Mp*PBD as did FP, with comparable K_d_ and EC_50_ values. Taken together, binding studies by FP and ITC have identified short peptidyl ligands with nano-molar affinities to *Mp*PBD (Figure 5E).

**Figure 5:**
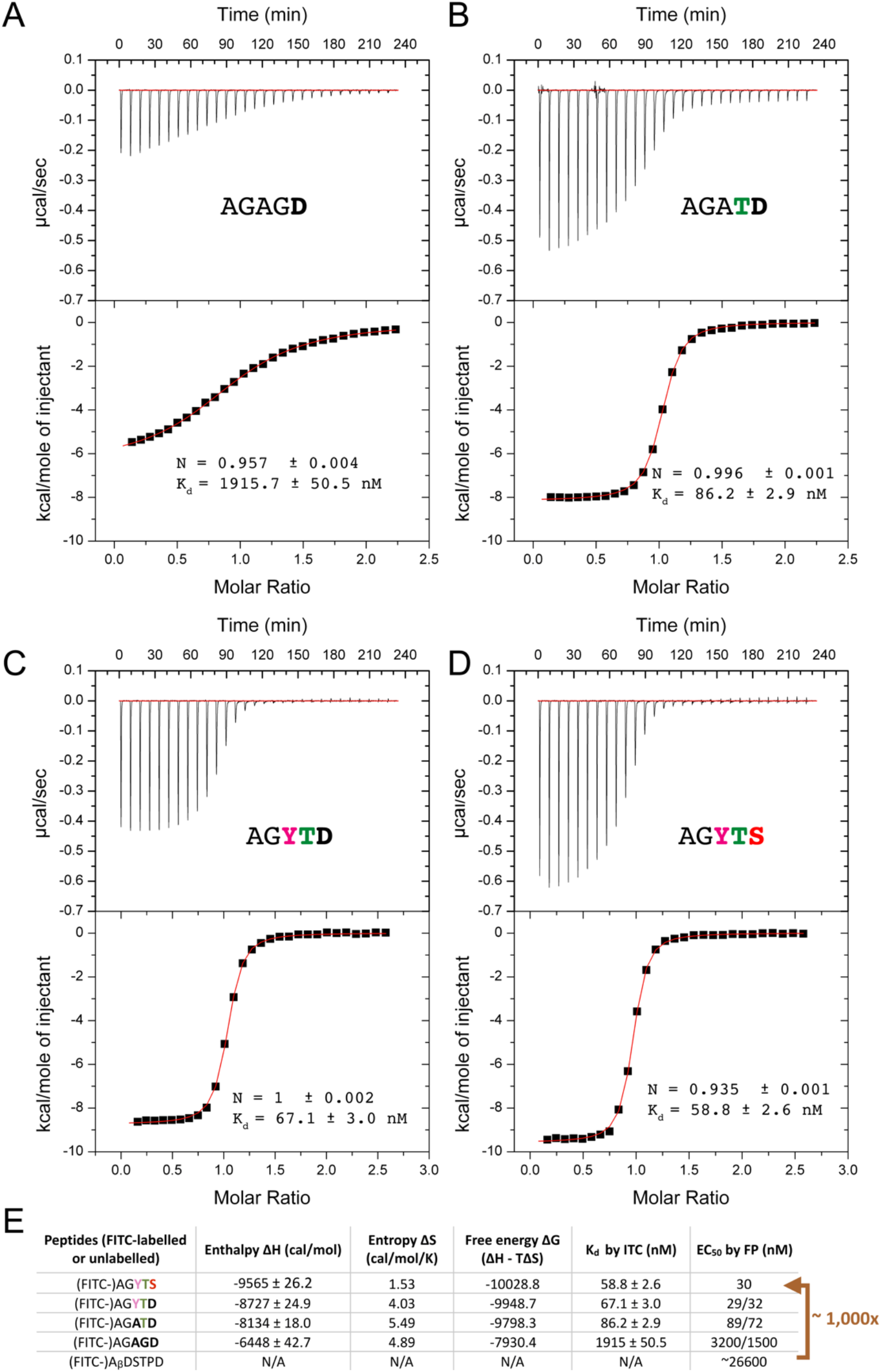
Isothermal titration calorimetry results of the binding of four unlabelled peptidyl ligands AGAGD (A), AGATD (B), AGYTD (C), and AGYTS (D) to *Mp*IBP_RIII1-4. **(E)** Table showing the thermodynamic parameters of the binding of the four unlabeled peptidyl ligands to *Mp*PBD calculated from ITC. K_d_ by ITC and EC_50_ by FP for the five representative peptidyl ligands of *Mp*PBD are shown. Red arrow compares the binding of the initially identified ligand that ends with “TPD” with the optimal sequences that end with “YTS” and “YTD” obtained from the structure-guided ligand optimization.

ITC indicated that significantly larger negative enthalpic (ΔH) contributions were involved in the binding of the potent ligands, AGATD, AGYTD, and AGYTS, to *Mp*PBD than for the moderate binder AGAGD (Figure 5E). In contrast, the positive entropic contributions (ΔS) were smaller for AGYTD and AGYTS when compared to AGAGD. The calculated thermodynamic profiles suggested that the enhancement of the peptide-protein interaction is primarily a result of a gain in polar interactions compared to a hydrophobic effect. This was supported by the structural comparison between the protein-peptide complex structures, which showed threonine or serine at peptide position 2 gained side-chain hydrogen bonding interactions with the protein compared to that of proline or glycine at the same position (Figure 3 vs Figure 1). However, additional high-resolution structural information was required to elucidate the basis for the stronger interactions between *Mp*PBD and its peptidyl ligands AGYTD and AGYTS.

### Molecular basis for potent binding by MpPBD ligands

To further reveal the molecular basis for the potent *Mp*PBD ligands, we solved X-ray crystal structures of *Mp*IBP_RIII1-4 in complex with the peptides AGYTS and AGYTD to a resolution of 1.8 Å (Supplementary file 1-Table S1, Figure 6, Supplementary file 1-Figure S6). Consistent with the other *Mp*PBD-peptide complexes, AGYTD and AGYTS had their terminal α-carboxyl group in contact with Ca1 and Ca2 via three ionic bonds with average lengths of approximately 2.5 Å (Figure 6, Figure 7A; Supplementary file 1-Table S4). The threonine residues in position 2 had their hydroxyl group bond with the side-chain amide of N333, as seen with FITC-A_β_DSTD and FITC-A_β_GPDSD (Figure 6). However, the YTD and YTS-containing peptides bound *Mp*PBD with more hydrogen bonds of shorter lengths on average compared to the weaker binders (Figure 7B; Supplementary file 1-Table S4), which is consistent with the large negative enthalpic contributions for the binding indicated by ITC. For example, the side chain of the terminal aspartate of AGYTD interacted with *Mp*IBP_RIII1-4 S295 and G296 with bond lengths of 2.5 Å and 2.8 Å, respectively (Figure 6A; Supplementary file 1-Table S4), while those at the same position for FITC-A_β_GPDSD and FITC-A_β_DSTD were at 3.2 Å and 3.5 Å, respectively (Figure 3). In contrast, the initially identified binding sequence that ends with TPD lacked these two bonds when bound by *Mp*PBD (Figure 1B).

**Figure 6:**
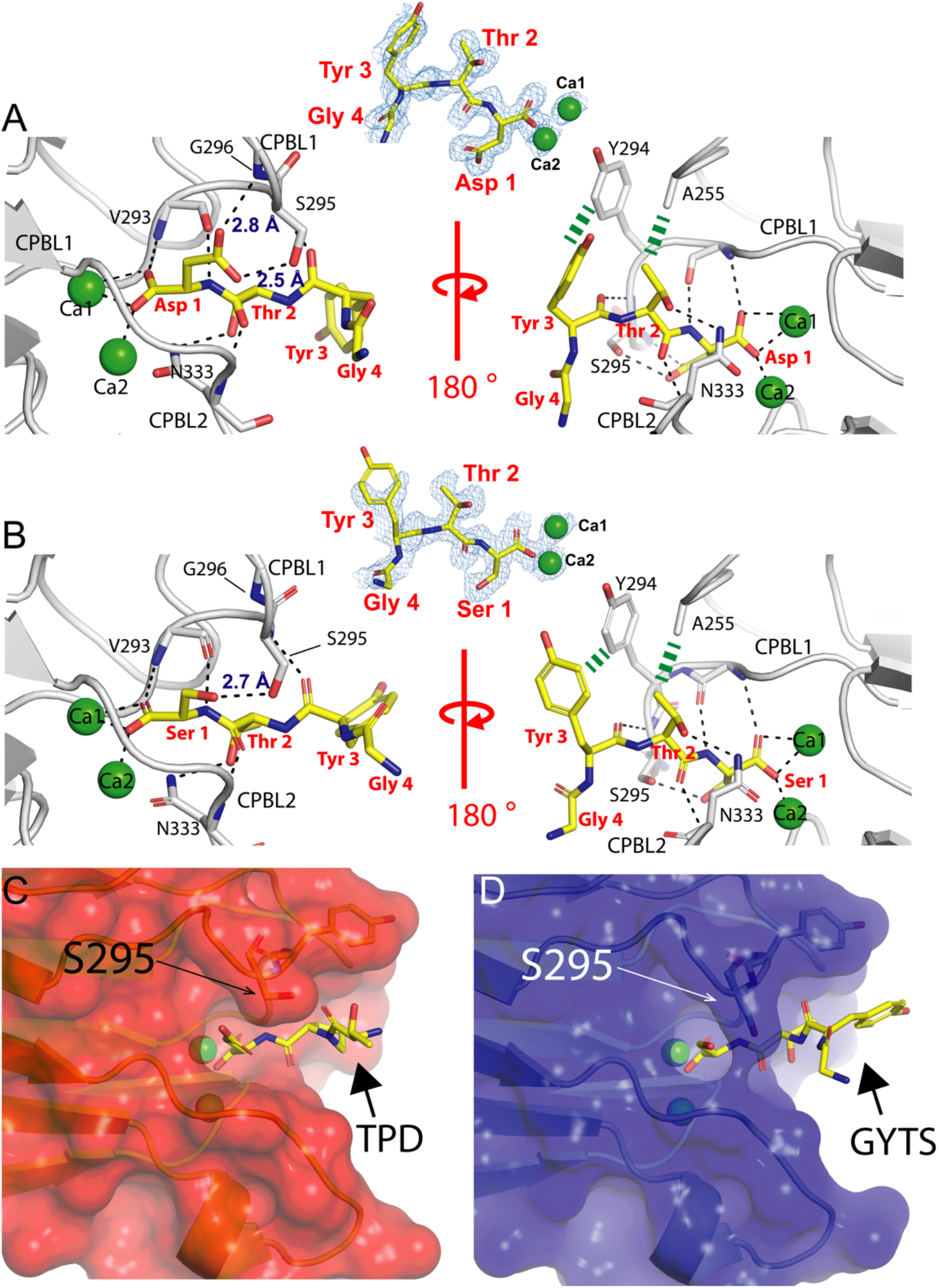
Atomic details of the interactions between *Mp*PBD and its nano-molar-affinity peptidyl ligands AGYTD (A) and AGYTS (B) revealed by X-ray crystallography. Color scheme is the same as in Figure 3. Thick green-dashed lines indicate hydrophobic contact between the protein and peptide tyrosine residues. The plasticity of the *Mp*PBD ligand-binding site is illustrated by their differences in complexing peptides that end with “TPD” (C) and “YTS” (D). Residue S295 that pointed to the side in C) and downward in (D) is indicated by black arrows. The protein is shown in surface representation while the peptide is shown in stick representation. CIF files can be found in Figure 6 – source data 2

**Figure 7:**
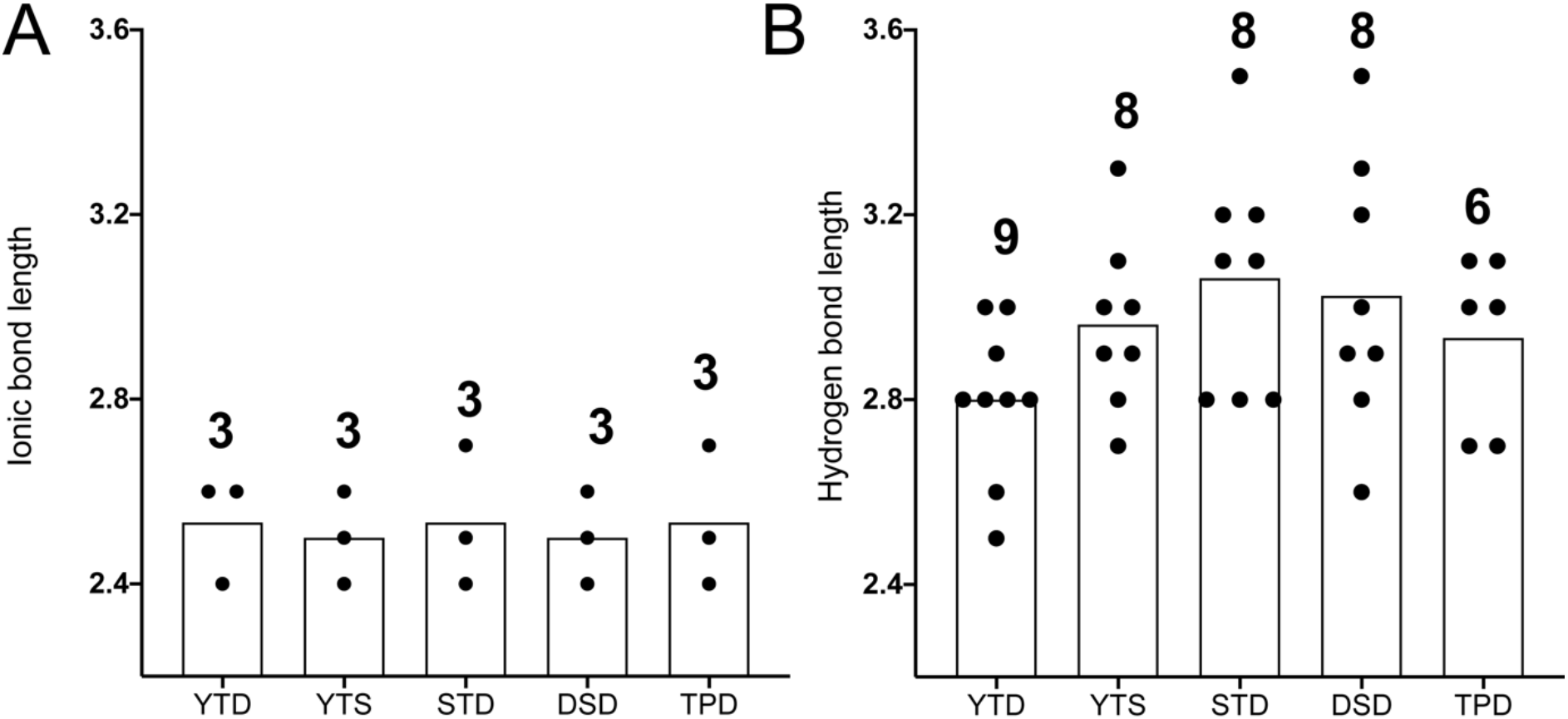
Length of the polar bonds between *MpPBD* and its peptidyl ligands revealed by X-ray crystal structures of protein-peptide complexes. Data bars are the average lengths for ionic (A) and hydrogen (B) bonds. Black data points indicate the length of individual bonds between the protein and peptide. Numbers above the data bars indicate the number of bonds at the protein-peptide interfaces. Details of the bonds are listed in Supplementary file 1-Table S4.

The tyrosine residues at position 3 of the YTD or YTS-containing peptides played a key role in their tight interactions with *Mp*PBD. Given the hydrophobic nature of its aryl side chain, the solvent-exposed peptide tyrosine appears to pack against Y294 of *Mp*PBD, which helps CPBLs to clench the peptide more tightly in the ligand-binding cavity (Figure 6A, B). Indeed, the hydrogen bond lengths between the peptide tyrosine main-chain oxygen (Tyr3-O) and protein S295 main-chain nitrogen (S295-N) is 2.9 Å for AGYTD and 3 Å for AGYTS, which are shorter than those at the same position for FITC-A_β_DSTD (3.2 Å) and FITC-A_β_GPDSD (3.3 Å) (Supplementary file 1-Table S4). These subtle structural differences underlined the molecular basis for the potent binding to *Mp*PBD by the peptides that end with YTD or YTS compared to the weaker ligands (Figure 6C, D).

Taken together, the presence of a tyrosine residue at position 3 of AGYTS and AGYTD stabilized the peptide-protein complexes via hydrophobic interactions. This corroborated with the results that other FITC-AGXTD peptides with aromatic residues at the 3rd position were also strong ligands for *Mp*PBD (Supplementary file 1-Table S3; Figure 4A). These atomic details explain why our structure-guided screening approach was extremely effective in obtaining potent ligands with nano-molar affinities for *Mp*PBD.

### AGYTD blocks MpPBD binding to diatoms

Having identified optimal peptidyl ligands for *Mp*PBD, we next tested their potential as antagonists to disrupt the PPI-mediated bacteria-diatom interactions that led to the discovery and characterization of this protein(27). Here we tested if AGYTD can block fluorescently labelled *Mp*IBP_RIII1-4 from binding to the Antarctic marine diatom *C. neogracile(44).* The porous silica cell wall (frustule) of *C. neogracile* is rectangular in shape with a length of roughly 10 μm and a width of 3-4 μm, with 1 −4 projections protruding from the corners (Figure 8A, Supplementary file 1-Figure S7). The binding of TRITC-labelled *Mp*IBP_RIII1-4 to *C. neogracile* resulted in red fluorescence evenly distributed around the cell centrally located inside of the diatom frustule (Figure 8B). At concentrations of 37.5 μM or above, AGYTD was extremely effective at blocking accumulation of the protein on the diatom, as the peptide outcompeted the cell surface proteins for binding *Mp*PBD and displaced ~ 95% of the fluorescent signal (Figure 8C, D, black bars). The inhibitory effect of AGYTD fell off by ~ 30% at 3.75 μM but was still more potent compared to the weaker *Mp*PBD ligand, AGAGD, even at a 100-fold higher concentration of 375 μM (Figure 8D, black and blue bars). While 37.5 μM AGAGD had a minimal effect on the binding of *Mp*PBD to diatoms, the potency of AGYTD diminished in the nano-molar concentrations. These results indicated that the effective inhibitory concentrations of the *Mp*PBD peptide ligands are significantly higher than the K_d_ values calculated for the protein-peptide interactions. This is likely due to the multi-valent effect originating from the presence of many nearby protein binding sites on the diatom cell membrane.

**Figure 8:**
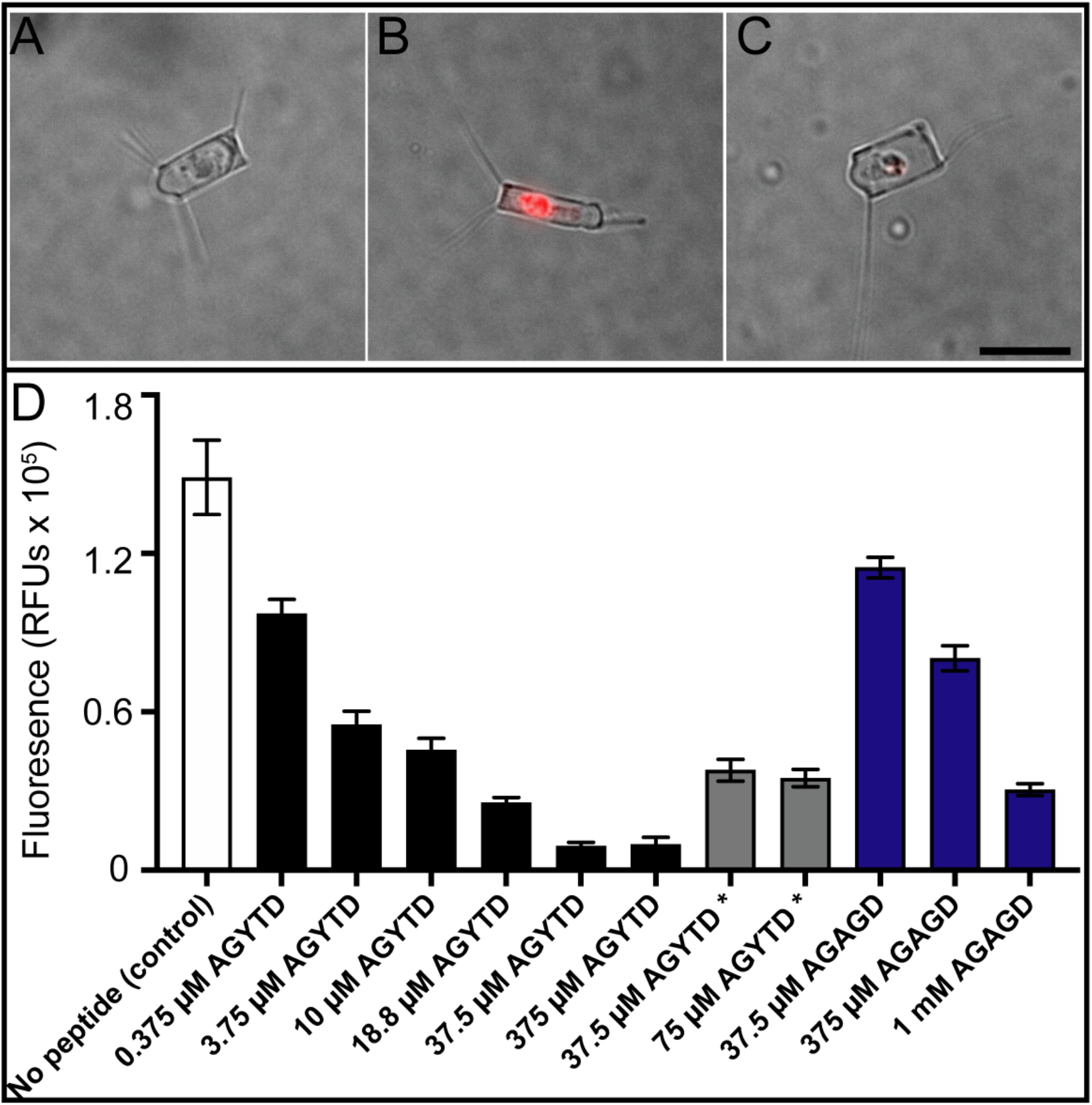
Inhibition of binding of *Mp*PBD to diatoms by competing peptides. Representative images showing: (A) an untreated *C. neogracile;* (B) a *C. neogracile* cell treated with 0.2 mg/mL TRITC-labelled *Mp*IBP_RIII1-4: (C) a *C. neogracile* cell treated with 0.2 mg/mL TRITC-labelled *Mp*IBP_RIII1-4 in the presence of 375 μM AGYTD. All three images (A) – (C) are at the same magnification with the black scale bar in (C) indicating 10 μm. (D) Fluorescence levels shown by the *C. neogracile* cells alone and those treated with TRITC-labelled *Mp*IBP_RIII1-4 and various concentrations of a strong or weak ligand, AGYTD and AGAGD, respectively. Each bar represents the quantification of average fluorescence from 30 individual diatoms. The white bar represents the control experiment where diatoms were treated with TRITC-labelled *Mp*IBP_RIII1-4 in the absence of peptide. Black bars represent experiments where diatoms were treated with 0.2 mg/mL TRITC-labelled *Mp*IBP_RIII1-4 and indicated concentrations of AGYTD. Grey bars with an asterisk underneath represent experiments where diatoms were incubated with 0.2 mg/mL TRITC-labelled *Mp*IBP_RIII1-4 before AGYTD was added. Blue bars represent experiments where diatoms were treated with 0.2 mg/mL TRITC-labelled *Mp*IBP RIII1-4 and indicated concentrations of AGAGD. Additional representative images are shown in Fig.S7.

Remarkably, AGYTD proved to be a potent antagonist that can displace *Mp*PBD prebound to the diatoms. The addition of 37.5 μM of AGYTD resulted in the dissociation of roughly 70% of the TRITC-labelled *Mp*IBP_RIII1-4 from the diatoms (Figure 8D, grey bars). Although the peptide was more effective at the same concentration when used as a prophylactic (Figure 8D, black bar), this important result demonstrated the potential of the peptide antagonists in disrupting pre-existing bacteria-host interactions, like those involved in biofilms.

## Conclusion and outlook

One major challenge in the field of PPI modulator discovery is a lack of starting points for the initial ligand identification. Here, a serendipitously discovered peptidyl ligand TPD has served as a lead for the development of a class of short peptides that bound *Mp*PBD with 1,000-fold higher affinity. This was largely achieved by systematically screening small libraries totaling 80 peptides that informed on the optimal residue in each of three terminal positions. Remarkably, these high-affinity ligands can serve as antagonists to disrupt *Mp*PBD from binding to the diatom cells that form symbiotic associations with the Antarctic bacterium.

Inhibition of pathogenic bacterial adhesion to human cells has yielded promising results in combating infections(17, 35, 38, 40, 45–47). The anti-adhesion strategy works by preventing and clearing the accumulation of bacteria at the sites of infections. In contrast to the conventional small-molecule antibiotics, the adhesin antagonists do not kill the bacteria, thus they are less likely to raise resistance. Furthermore, given that adhesins are localized to the cell surfaces, their antagonists are not required to penetrate or be taken into the cells, simplifying their applications. These key attributes justified the pursuit of novel adhesin modulators to disrupt PPI-mediated bacterial adhesion to hosts to treat bacterial infections. Homologs of *Mp*PBD with a conserved ligand-binding site are found in human pathogens such as *Vibrio cholerae*(27, 28, 48), the causative agent of cholera and others such as *Aeromonas veronii* that cause infections ranging from diarrhoea to wound infections and sepsis(49, 50), and the flesh-eating bacterium *Vibrio vulnificus(51)* (Figure S8). Therefore, there is an opportunity to use this conceptual approach to devise antagonists to disrupt the PPIs involved in bacterial infections. With the emergence of antibiotic-resistant pathogens, this work gives insight into how microbes might be controlled through the modulation of PPI-related adhesion to their hosts. In future research, it will be of interest to design adhesin modulators consisting of D-peptides, which are non-metabolizable and are thus resistant to proteolytic degradation, as well as cyclic peptides that might achieve higher affinity via favourable entropic contributions.

## Materials and Methods

### Peptide synthesis

FITC-labelled peptides FITC-A_β_GPDSD, FITC-A_β_DSTPD, FITC-A_β_DSTD and FITC-AβGPDD were synthesized by Fmoc solid-phase peptide synthesis, either manually or using an automated Intavis MultiPep RSi peptide synthesizer. The protected amino acids (linked to Wang resins) and FITC were purchased from Novabiochem and Sigma-Aldrich. Crude peptides were then analyzed and purified by high-pressure liquid chromatography (HPLC) using a preparative reversed-phase column with MS detection. The peptides were freeze-dried and stored at – 30 °C. All other FITC-labelled peptides used in the structure-guided ligand optimization procedures by fluorescence polarization (FP) and the four unlabeled peptides (AGAGD, AGYTD, ATYTS and AGATD) used in the ITC measurements and X-ray crystallography were purchased from GenicBio (Shanghai, China). HPLC and MS spectra for representative peptides are shown in Supplementary file 1-Figure S9-S21.

### Fluorescence polarization

The FITC-labelled peptides were dissolved in FP buffer (10 mM HEPES, pH 7.4, 150 mM NaCl, 0.1% Tween 20, 1 mg/mL BSA) to a final concentration of 10 nM. Dilution series of *Mp*IBP_RIII1-4 were made on round-bottom 384-well plates (Corning, Black). The protein-peptide mixture in the plate was incubated at room temperature for at least 30 min before the fluorescence polarization was measured using a Tecan Infinite F500 plate reader (excitation 485 nm, emission 535nm). All measurements were performed in triplicate, and the data were plotted with the GraphPad Prism 8 software using a non-linear regression analysis method (single-site binding model).

### Co-crystallization, X-ray diffraction and structure solutions of *Mp*PA14 with peptide ligands

The original *Mp*IBP_RIII1-4 construct self-associated in the crystal by inserting its C-terminal “TPD” sequence into the ligand-binding site of a symmetry-related molecule. This crystal contact competed with free peptides for binding *Mp*PBD and interfered with the crystallization of peptide-protein complexes. Thus, *Mp*IBP_RIII1-4 protein used for co-crystallography was truncated by 12 amino acids from the original construct, which ended at the residue N507 instead of D519. In all other respects *Mp*IBP_RIII1-4 was produced, purified, and crystallized as previously described(27, 52). Co-crystallization of *Mp*IBP_RIII1-4 with various peptides was performed using the “microbatch-under-oil” method by mixing equal volumes of ~ 5 mg/mL protein with a precipitant solution composed of approximately 0.1 M calcium chloride, 0.1 M sodium acetate (pH 4.6), 30% (w/v) PEG400 and 1-2 mM of different peptides, including FITC-A_β_GPDSD, FITC-A_β_DSTD, and the unlabeled peptides AGYTD and AGYTS.

X-ray crystallographic data were collected either at the P11 beamline of the PETRA III facility at DESY (Hamburg, Germany) or at the 08ID-1 beamline of the Canadian Light Source synchrotron facility via remote access. Data were indexed and integrated with X-ray Detector Software (XDS)(53) and CCP4-Aimless(54) or the DIALS/xia2(55, 56) in the CCP4i2 software suite. The structure solutions for all complexes were obtained by using molecular replacement using the *Mp*IBP_RIII1-4 structure (PBD: 5K8G) as the search model(27). The structures were refined using CCP4-Refmac5(57).

### Isothermal Titration Calorimetry

Isothermal calorimetric titration (ITC) measurements were performed at 30 C using a MicroCal VP-ITC instrument (Malvern). *Mp*IBP_RIII1-4 was dialyzed overnight in a buffer of 50 mM Tris-HCl, pH 9, 150 mM NaCl, 5 mM CaCl_2_. Next, the protein was diluted to approximately 20 μM and was mixed with serial 5-μl aliquots of 200 μM of each of the four peptide solutions (AGAGD, AGATD, AGYTD and AGYTS). Peptide solutions were automatically added by a rotating syringe (400 RPM) at 5-min intervals into the *Mp*IBP_RIII1-4 solution for a total of 50 injections. The data were analyzed by Origin software Version 5.0 (MicroCal).

### Diatom binding experiments

The Antarctic diatom, *Chaetoceros neogracile,* was cultured as previously described(27, 44). TRITC-labelled *Mp*IBP_RIII1-4 (TRITC-*Mp*IBP_RIII1-4, 0.2 mg/mL) in the presence or absence of peptides was incubated with diatoms in buffer (50 mM Tris-HCl pH 9, 300 mM NaCl, 5 mM CaCl_2_) with gentle mixing for 2 h. Next, diatoms were pelleted by centrifugation for 3 min at 4,500 x g, and the resulting supernatant was discarded. This procedure was repeated three times to wash out unbound TRITC-*Mp*IBP_RIII1-4 before the diatom pellet was finally resuspended in 20 μL of buffer, which was then examined on slides by fluorescence microscopy. In parallel experiments to test if the strong ligand AGYTD could compete off the TRITC-*Mp*IBP_RIII1-4 that was already bound to diatoms, TRITC-*Mp*IBP_RIII1-4 was incubated with diatom for 1.5 h before the peptide AGYTD was added. The remainder of the experiment followed the same procedure as described above.

Images were obtained using an Olympus IX83 inverted fluorescence microscope equipped with an Andor Zyla 4.2 Plus camera. Quantification of the fluorescence intensity was done using Fiji ImageJ. The corrected total cell fluorescence (CTCF) was calculated using the formula: CTCF = Integrated Density – (Area of selected cell x Mean Fluorescence of the background)(58). Quantification of 30 individual diatom cells was done for each treatment. As diatom cell aggregates produce overexposed fluorescence while those cells lacking a silica frustule have damaged plasma membranes necessary for protein binding, they were excluded from the measurements. Graphs were made using Graphpad Prism.

## Data availability

X-ray crystal structure coordinates solved in this study have been deposited in the Protein Data Bank with accession codes of 6X6Q (*Mp*IBP_RIII1-4 – FITC-AβGPDSD), 6X6M (*Mp*IBP_RIII1-4 – FITC-A_β_DSTD), 6X5W (*Mp*IBP_RIII1-4 – AGYTD), and 6X5V (*Mp*IBP_RIII1-4 – AGYTS). The data that support the findings of this study are available from the corresponding author P.L.D upon reasonable request.

## Acknowledgements

We thank Dr. John Allingham for access to his home source X-ray diffractometer at Queen’s University, and to staff members at the Canadian Light Source in Saskatoon, Canada and the PETRA III facility at DESY in Hamburg, Germany for access to data collection at these synchrotrons. We are grateful to Dr. EonSeon Jin, Hanyang University, Seoul, for the gift of the diatom, *Chaetoceros neogracile* and Dr. Saeed Rismani Yazdi for assistance with the diatom cultures. We are indebted to Mr. Kim Munro at the Protein Function Discovery unit at Queen’s University for his assistance with acquiring and interpreting ITC data. We thank Ms. Irene van Oekel for preliminary tests on the binding of peptidyl ligands to *Mp*PBD and Mr. Joost van Dongen for analytical support. This project was funded by a Natural Science and Engineering Research Council (NSERC, http://www.nserc-crsng.gc.ca/index_eng.asp) Discovery Grant (RGPIN-2016-04810) to PLD who holds the Canadian Research Chair in Protein Engineering. IKV acknowledges financial support by the European Union (ERC-2014-StG Contract No.635928) and the Dutch Science Foundation (NWO ECHO Grant No. 712.016.002).

## Author contributions

S.G. and P.L.D. conceived the study, designed the experiments, and wrote the manuscript. S.G. performed co-crystallization, data collection, and structure determination of the X-ray crystal structures. S.G. performed FP binding experiments and analyzed data with assistance of D.C.S.. L.G.M. and D.C.S. performed peptide synthesis and purification. H.Z. and C.S. performed the diatom binding experiments and analyzed data. C.O., L.B., and I.K.V. provided critical feedback to S.G. throughout the project and contributed to the critical editing of drafts of the manuscript.

