## Supplementary material for "Molecular basis for inhibition of adhesin-mediated bacterial-host interactions through a novel peptide-binding domain": SI

### For

**Table of contents:**

Figure S1. Amino-acid sequence and secondary structural elements of *Mp*IBP_RIII1-4.

Figure S2-5. FP assays of the interaction between *Mp*IBP_RIII1-4 and 80 FITC-labeled peptides during the four rounds of screening.

Figure S6. Ligand-binding site of *Mp*PBD in complex with various peptides.

Figure S7. Representative images of diatoms with various TRITC-*Mp*PBD and peptides treatments shown in Figure 8.

Figure S8. Amino-acid alignment of *Mp*PBD with PBDs from pathogenic bacteria including *Shewanella putrefaciens, Aeromonas veronii, Shewanella oneidensis, Vibrio vulnificus* and *Vibrio cholerae.*

Figure S9-21. Representative analytical LC/MS of the purified FITC-labelled and unlabelled peptides used in this study.

Table S1. Statistics for the crystallographic data of *Mp*PBD in complex with four peptide ligands.

Table S2-3: Average $EC_{50}$ values calculated from the binding of FITC-labelled peptides to *Mp*PBD determined by FP.

Table S4. Ionic and hydrogen bonds at the protein-peptide interfaces. Length of the bonds are indicated.

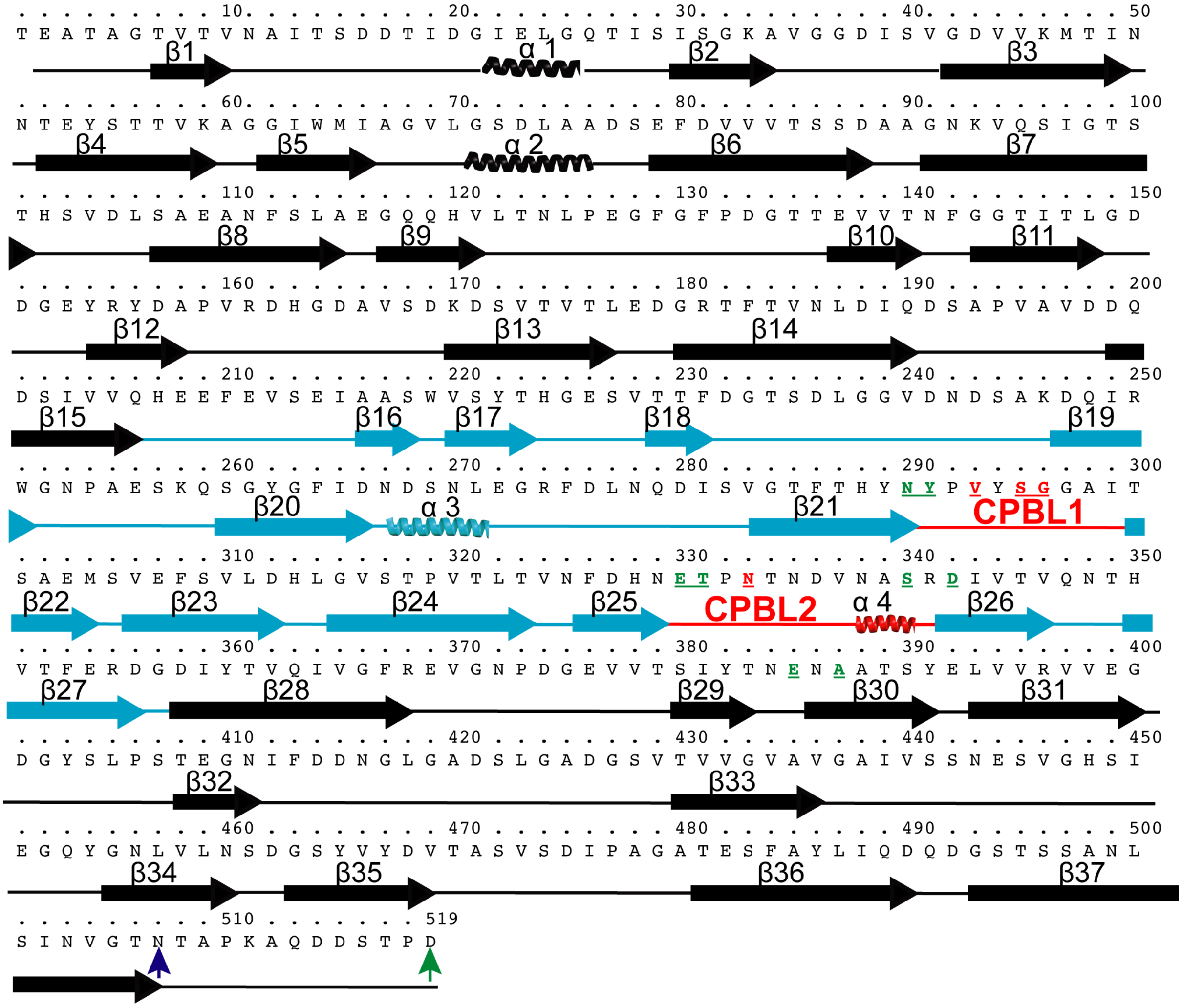


**Figure S1. Amino-acid sequence and secondary structural elements of *Mp*IBP_RIII1-4.** Secondary structural elements for *Mp*PBD are colored cyan. The Calcium-Peptide-Binding Loops (CPBLs) 1 and 2 are indicated and are colored red. Residues that hydrogen bond with the peptide are bolded and underlined in red, while those coordinating Ca1 and Ca2 are colored green. The ending residue D519 for the original construct used in a previous work is indicated with a green arrowhead, while the C-terminal residue N507 for the new construct used in this study is marked by a blue arrowhead.

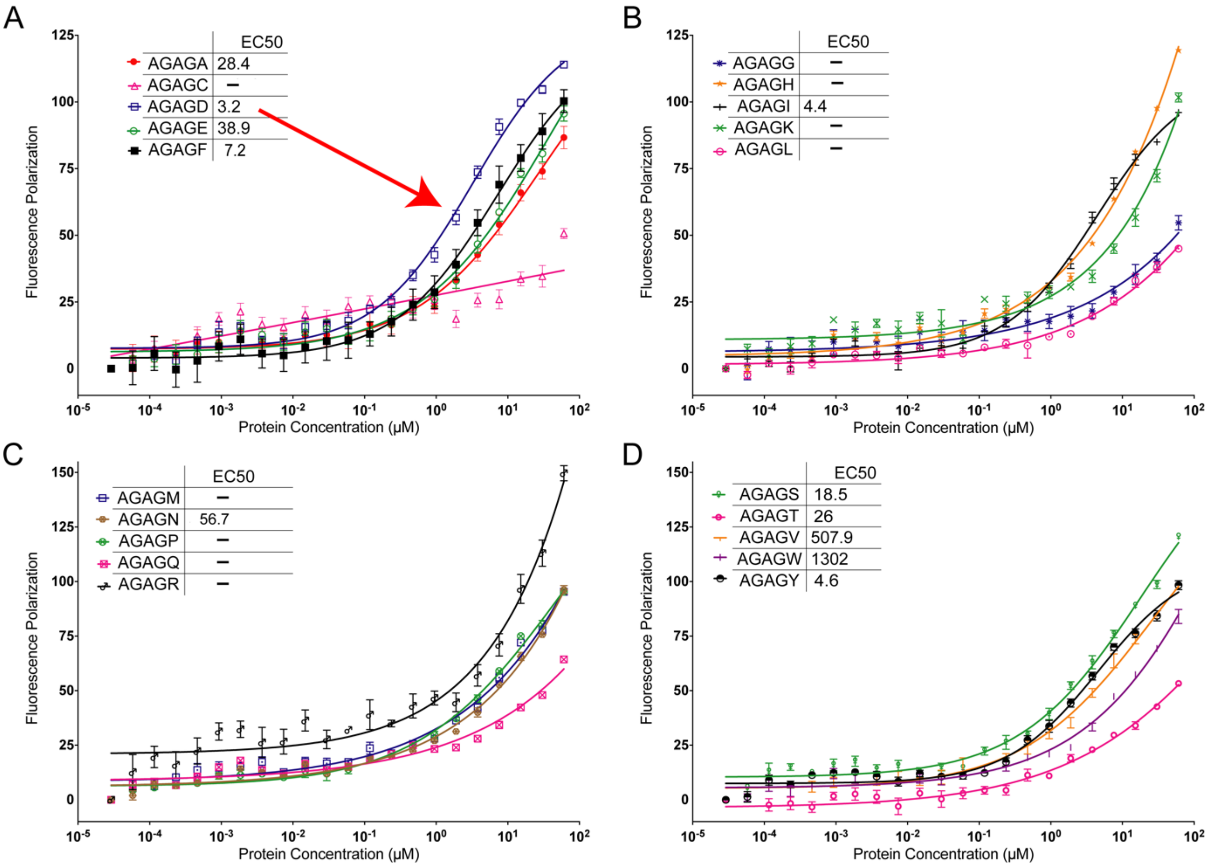


**Figure S2. FP assays of the interaction between *Mp*IBP_RIII1-4 and 20 FITC-labeled peptides AGAGX.** Each plot shows results of five different peptides. Red arrow points to the results of FITC-AGAGD binding to *Mp*IBPRIII_1-4, which is the strongest ligand of the 20 AGAGX peptides. (A) FP assays of the binding of five peptides from FITC-AGAGA to FITC-AGAGF to *Mp*IBP_RIII1-4. (B) FP assays of the binding of five peptides from FITC-AGAGG to FITC-AGAGL to *Mp*IBP_RIII1-4. (C) FP assays of the binding of five peptides from FITC-AGAGM to FITC-AGAGR to *Mp*IBP_RIII1-4. (D) FP assays of the binding of five peptides from FITC-AGAGS to FITC-AGAGY to *Mp*IBP_RIII1-4. All experiments were performed in triplicate. Color and symbol coding for each listed peptide within the set is show in the key for each quadrant of the figure, alongside their $E_{50}$ values.

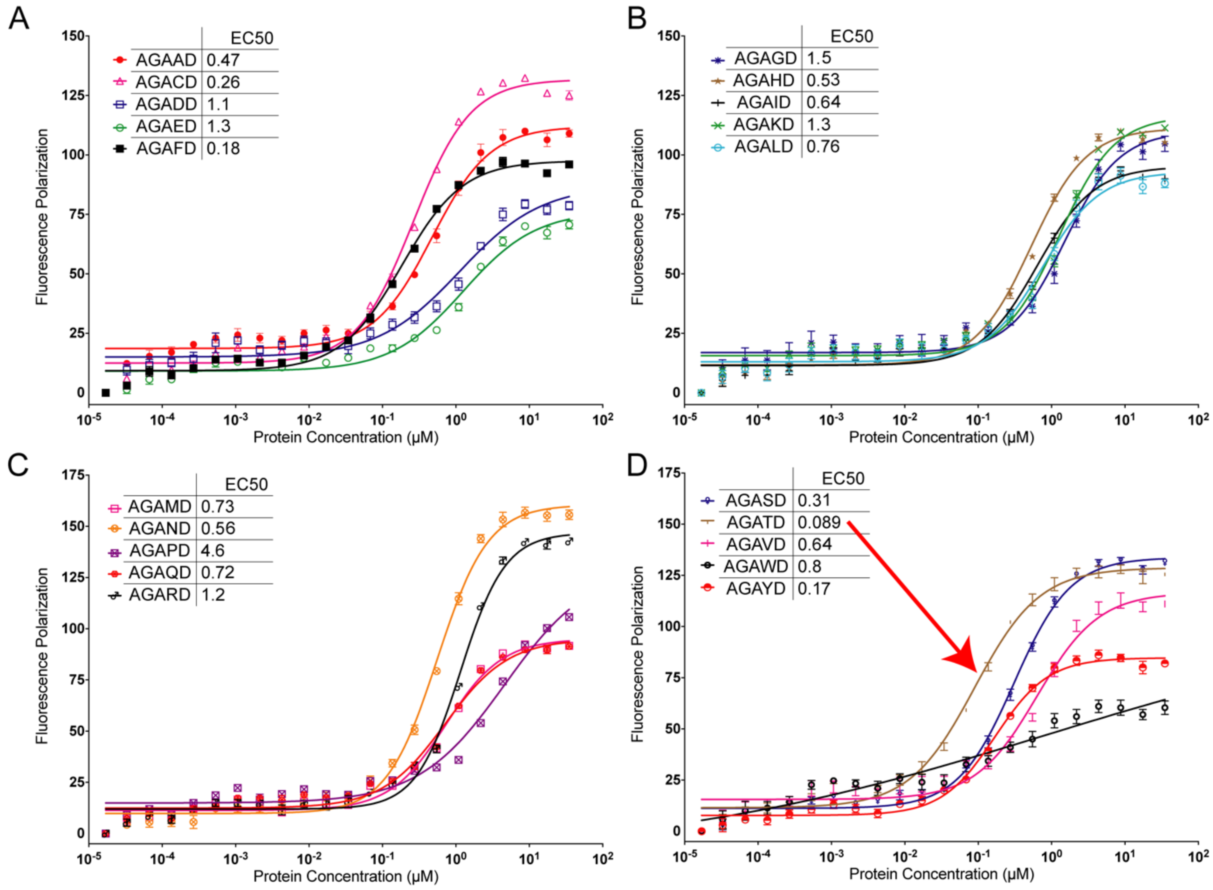


**Figure S3. FP assays of the interaction between *Mp*IBP_RIII1-4 and 20 FITC-labeled peptides AGAXD.** Each plot shows results for five of the 20 FITC-AGAXD peptides. Red arrow points to the results of FITC-AGATD (D) binding to *Mp*IBPRIII_1-4, which is the strongest ligand among the 20 AGAXD peptides. (A) FP assays of the binding of five peptides from FITC-AGAAD to FITC-AGAFD to *Mp*IBP_RIII1-4. (B) FP assays of the binding of five peptides from FITC-AGAGD to FITC-AGALD to *Mp*IBP_RIII1-4. (C) FP assays of the binding of five peptides from FITC-AGAMD to FITC-AGARD to *Mp*IBP_RIII1-4. (D) FP assays of the binding of five peptides from FITC-AGASD to FITC-AGAYD to *Mp*IBP_RIII1-4. All experiments were performed in triplicate. Color and symbol coding for each listed peptide within the set is show in the key for each quadrant of the figure, alongside their $E_{50}$ values.

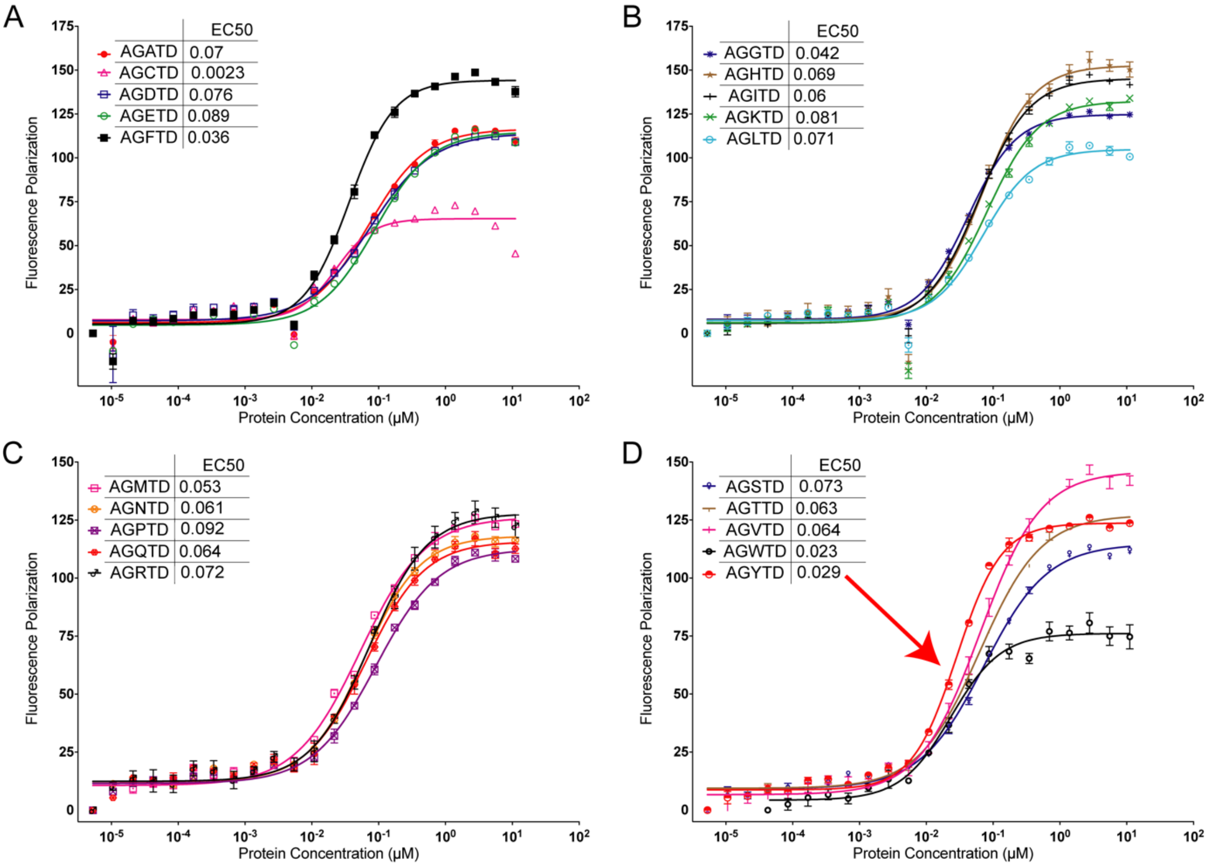


**Figure S4. FP assays of the interaction between *Mp*IBP_RIII1-4 and 20 FITC-labeled peptides AGXTD. Each plot shows results of five different peptides.** Red arrow points to the results of FITC-AGYTD binding to *Mp*IBPRIII_1-4, which is the strongest ligand of the 20 AGXTD peptides. (A) FP assays of the binding of five peptides from FITC-AGATD to FITC-AGFTD to *Mp*IBP_RIII1-4. (B) FP assays of the binding of five peptides from FITC-AGGTD to FITC-AGLTD to *Mp*IBP_RIII1-4. (C) FP assays of the binding of five peptides from FITC-AGMTD to FITC-AGRTD to *Mp*IBP_RIII1-4. (D) FP assays of the binding of five peptides from FITC-AGSTD to FITC-AGYTD to *Mp*IBP_RIII1-4. All experiments were performed in triplicate. Color and symbol coding for each listed peptide within the set is show in the key for each quadrant of the figure, alongside their $E_{50}$ values.

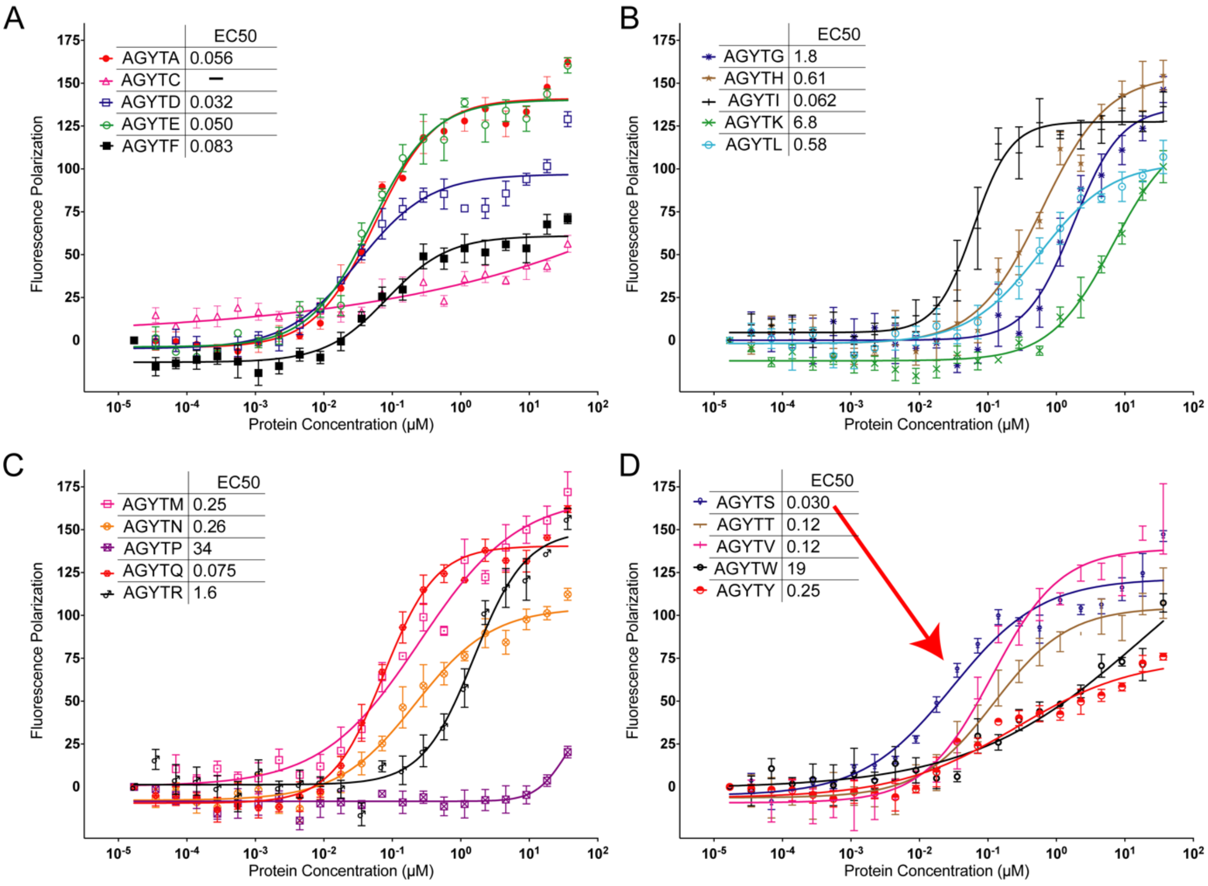


**Figure S5. FP assays of the interaction between *Mp*IBP_RIII1-4 and 20 FITC-labeled peptides AGYTX.** Each plot shows results of five different peptides. Red arrow points to the results of FITC-AGYTS binding to *Mp*IBPRIII_1-4, which is the strongest ligand of the 20 AGXTD peptides. (A) FP assays of the binding of five peptides from FITC-AGYTA to FITC-AGYTF to *Mp*IBP_RIII1-4. (B) FP assays of the binding of five peptides from FITC-AGYTG to FITC-AGYTL to *Mp*IBP_RIII1-4. (C) FP assays of the binding of five peptides from FITC-AGYTM to FITC-AGYTR to *Mp*IBP_RIII1-4. (D) FP assays of the binding of five peptides from FITC-AGYTS to FITC-AGYTY to *Mp*IBP_RIII1-4. All experiments were performed in triplicate. Color and symbol coding for each listed peptide within the set is show in the key for each quadrant of the figure, alongside their $E_{50}$ values.

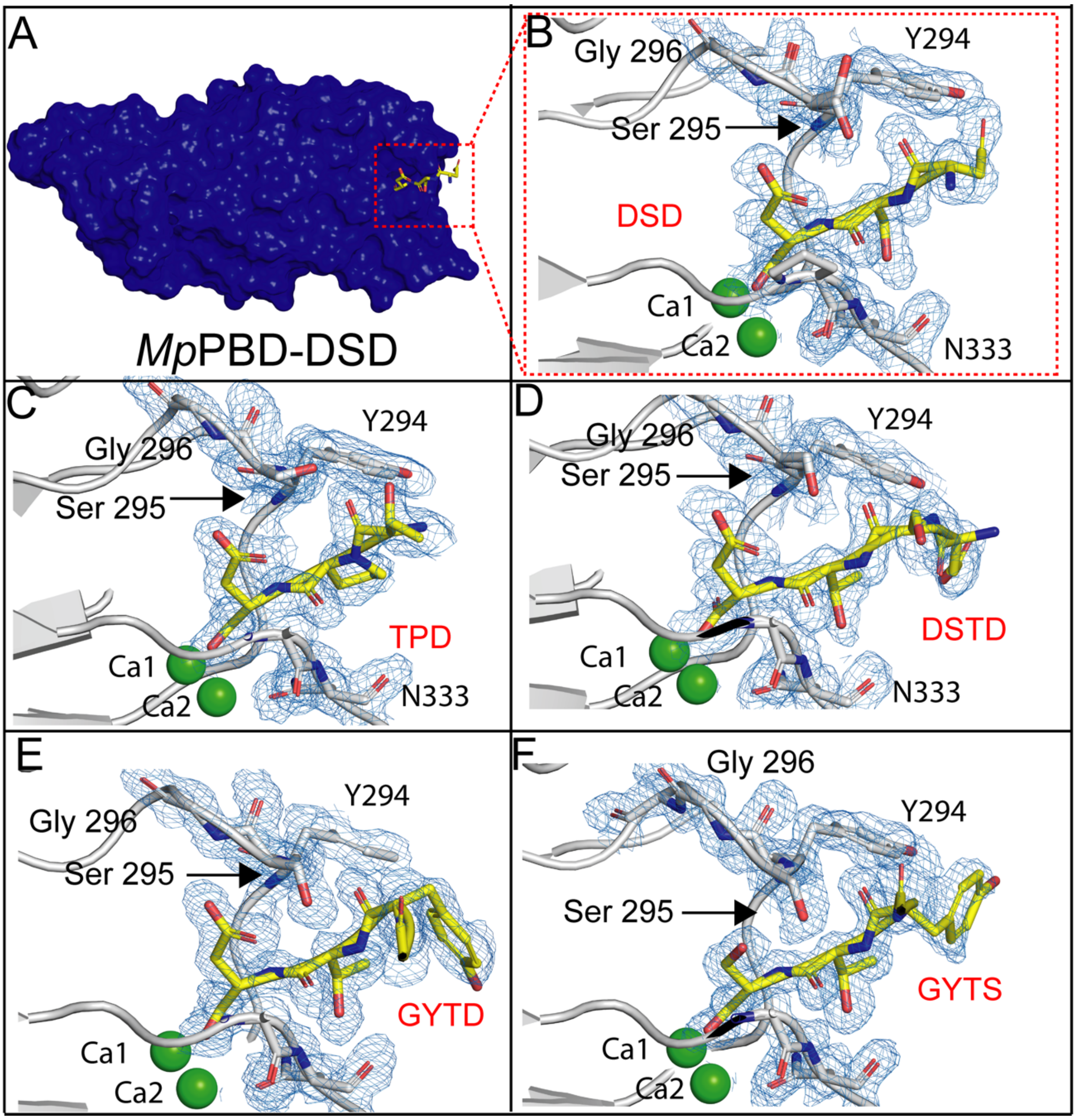


**Figure S6. Ligand-binding site of *Mp*PBD in complex with various peptides.** A) surface representation of *Mp*PBD bound by peptide "DSD" (stick representation). Zoomed-in views of the *Mp*PBD ligand-binding site in complex with peptidyl sequences of "DSD" (B), "TPD" (C), "DSTD" (D), "GYTD" (E), "GYTS" (F). Electron density maps (2 $F_o - F_c$, σ =1) of *Mp*IBP_RIII1-4 residues Y294-G296, and N333 in the CPBLs are shown as blue meshes. Carbon atoms of the peptides are colored in yellow while those for the protein are colored in grey. Oxygen atoms are red, nitrogen atoms are blue, while $Ca^{2+}$ ions are shown as green spheres. Amino acids involved in protein-peptide interactions are shown in stick representation.

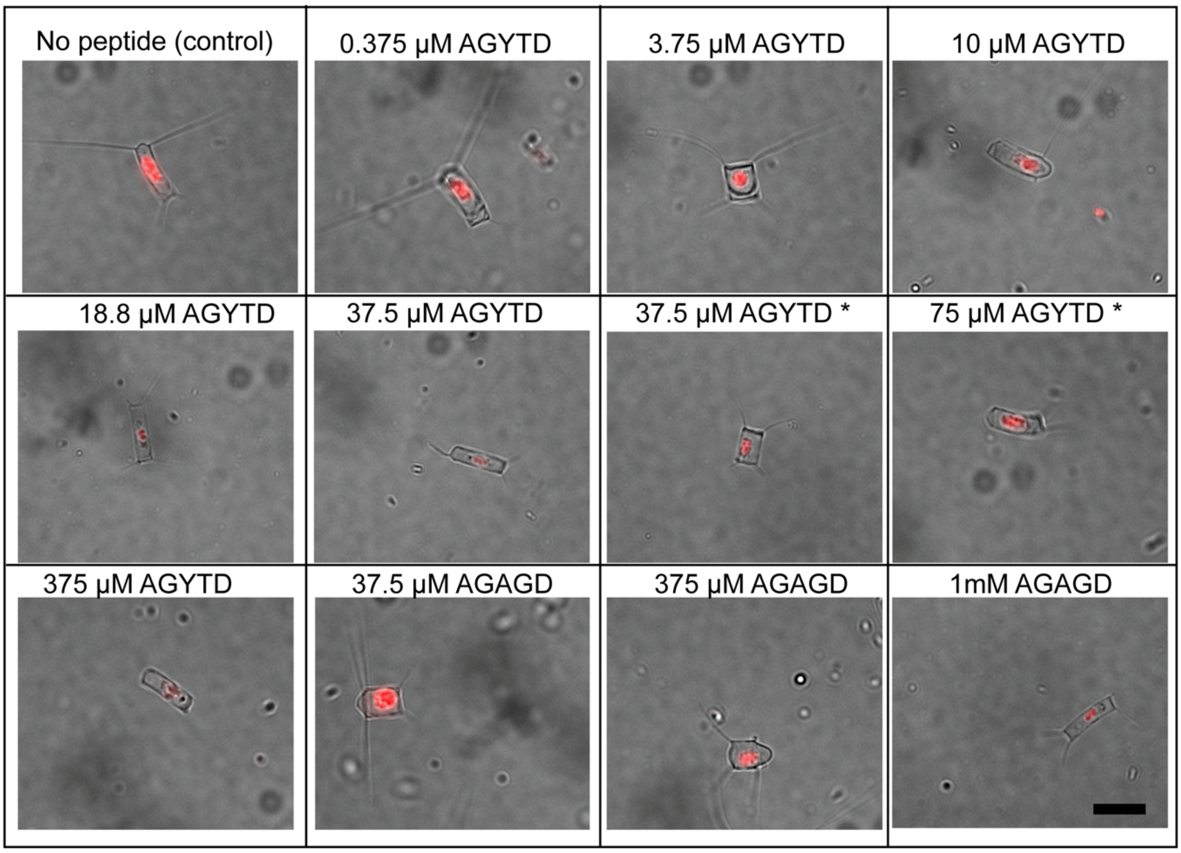


**Figure S7. Representative images of diatoms with various TRITC-*Mp*PBD and peptides treatments shown in Figure 8.** Asterisks (*) represent experiments where diatoms were incubated with TRITC-*Mp*PBD before AGYTD was added. Scale bar represents 10 µm.

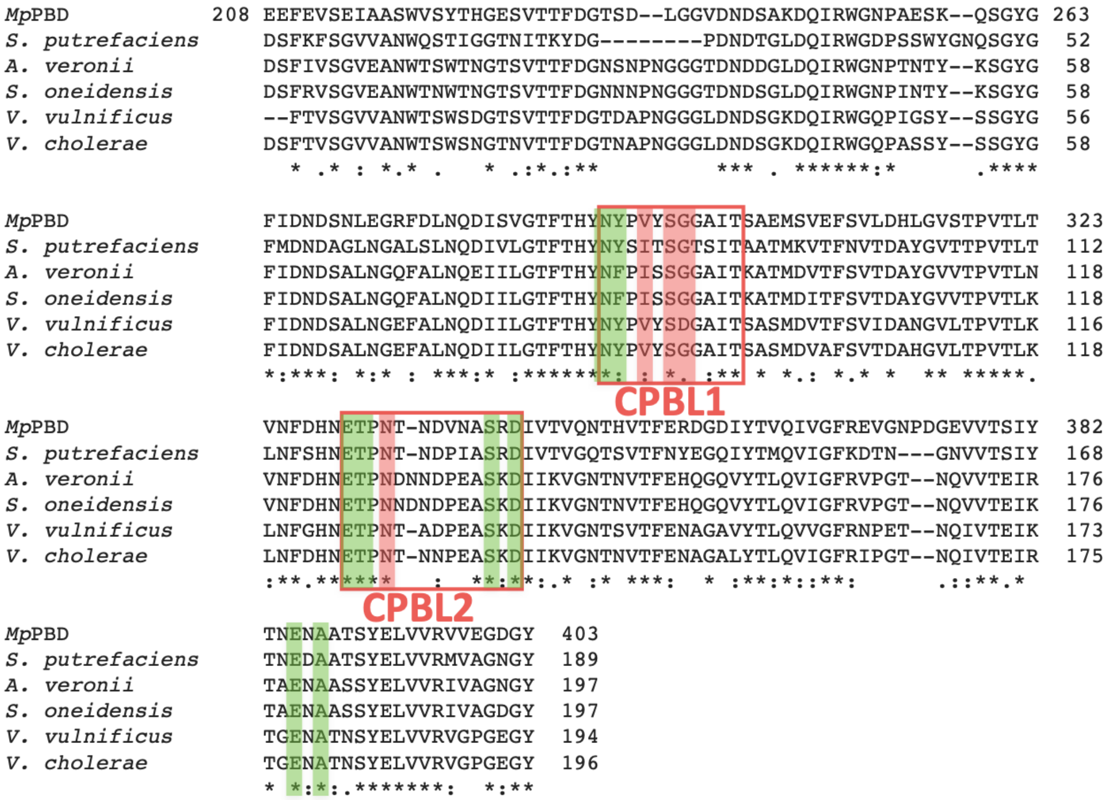


**Figure S8. Amino-acid alignment of *Mp*PBD with PBDs from pathogenic bacteria including *Shewanella putrefaciens, Aeromonas veronii, Shewanella oneidensis, Vibrio vulnificus* and *Vibrio cholerae.*** Residue similarity is indicated as follows; match (*), high similarity (:), low similarity (.). Residues that constitute CPBLs 1 and 2 are indicated with red boxes. Residues that coordinate Ca1 and Ca2 are indicated with green shades, while those that interact with peptides are shaded red. *Mp*PBD was numbered in the same way as shown in Figure S1.

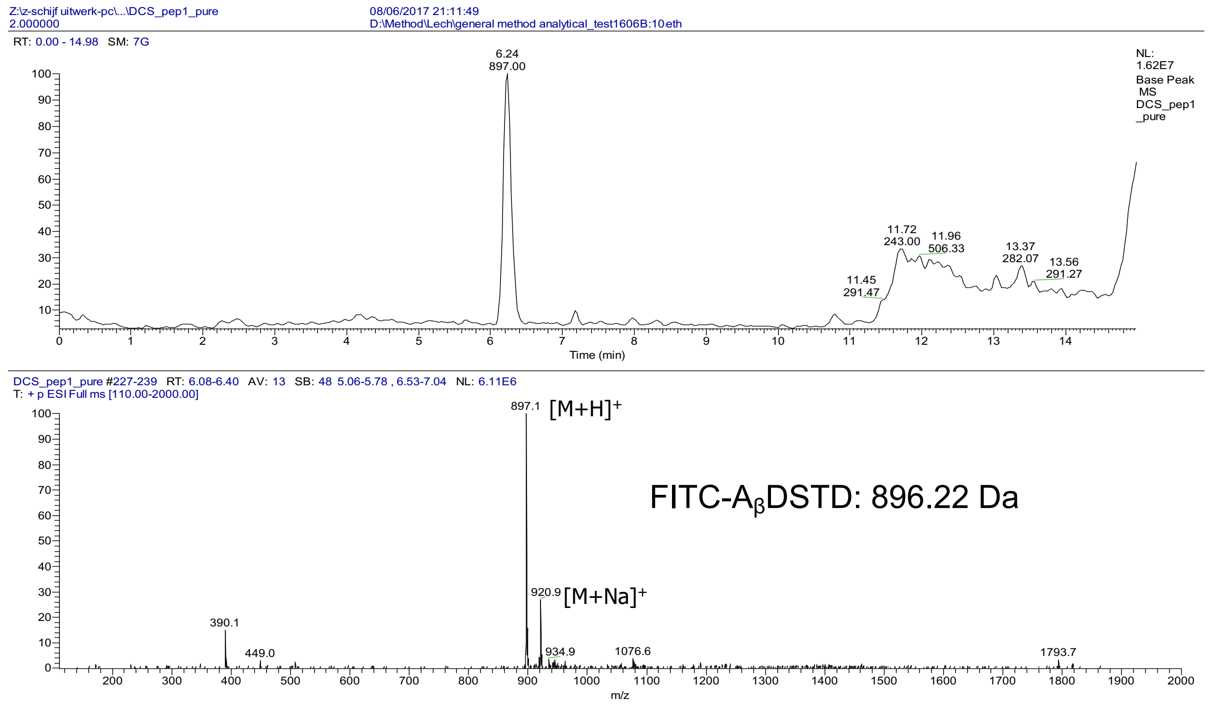


**Figure S9. Analytical LC/MS of the purified FITC-labelled peptide A$_β$DSTD.** The graphs represent the base peak chromatogram (upper panel) and the mass spectrum (lower panel).

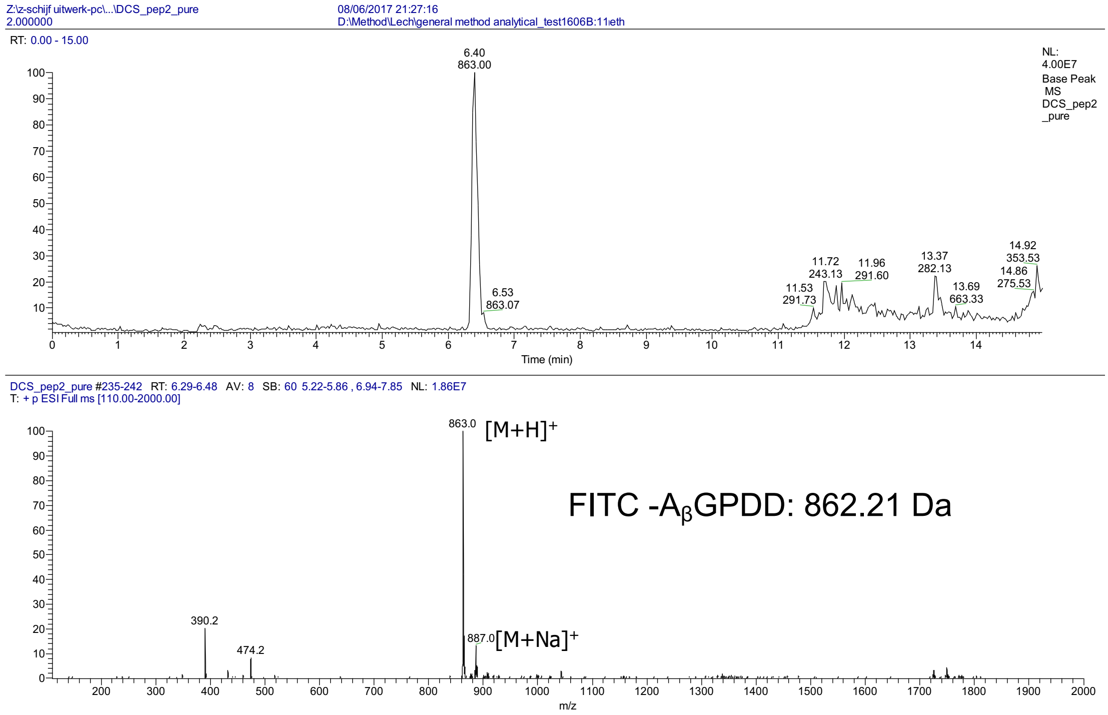


**Figure S10. Analytical LC/MS of the purified FITC-labelled peptide A$_β$GPDD.** The graphs represent the base peak chromatogram (upper panel) and the mass spectrum (lower panel).

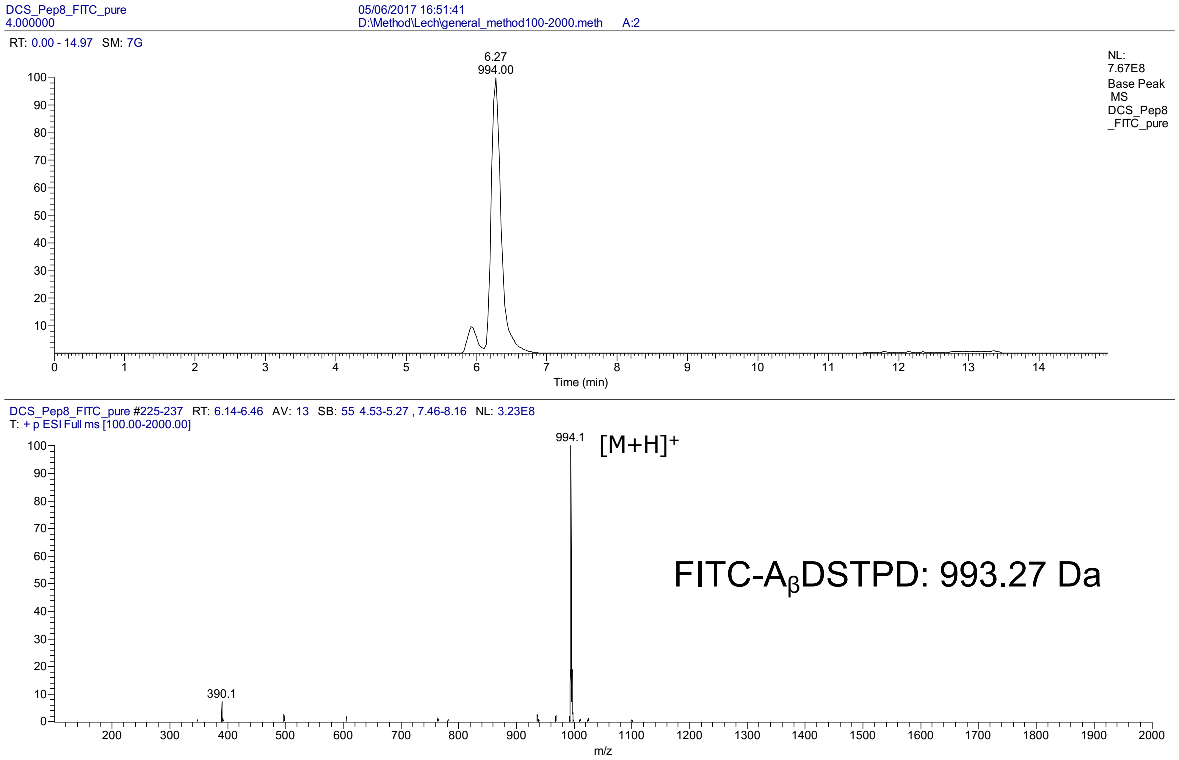


**Figure S11. Analytical LC/MS of the purified FITC-labelled peptide $A_\beta$DSTPD.** The graphs represent the base peak chromatogram (upper panel) and the mass spectrum (lower panel).

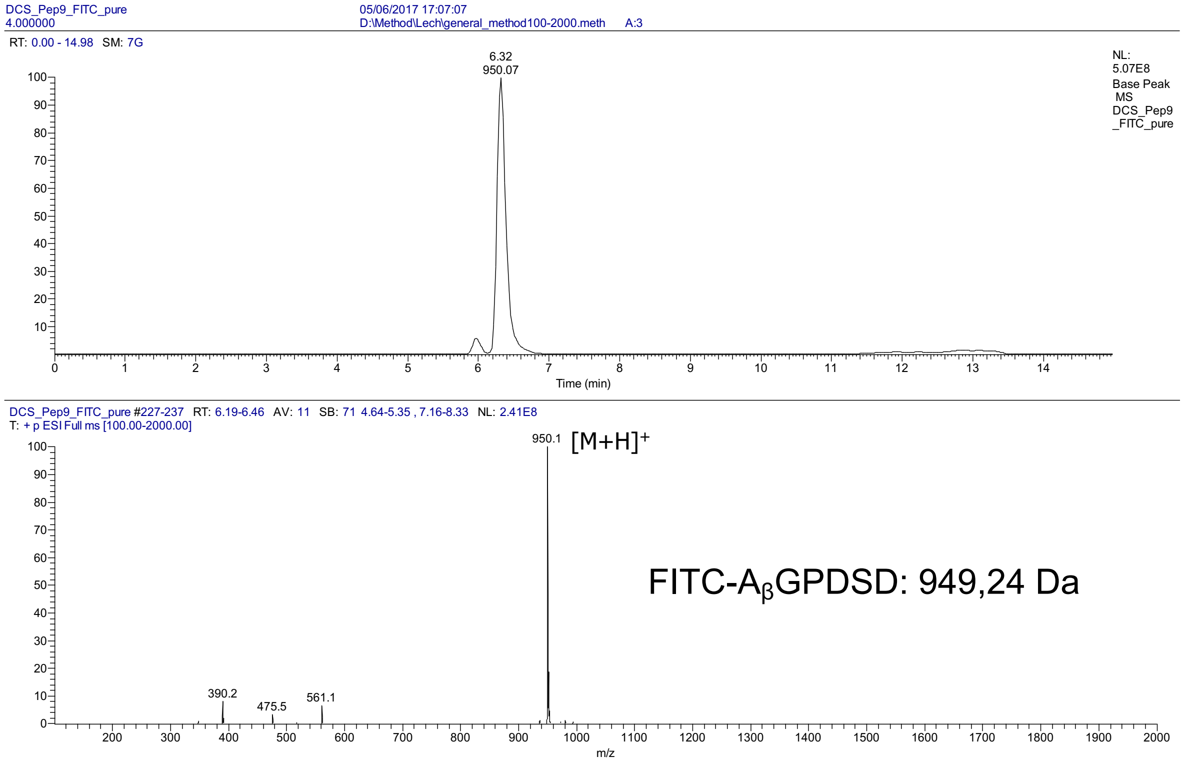


**Figure S12. Analytical LC/MS of the purified FITC-labelled peptide $A_\beta$GPDSD.** The graphs represent the base peak chromatogram (upper panel) and the mass spectrum (lower panel).

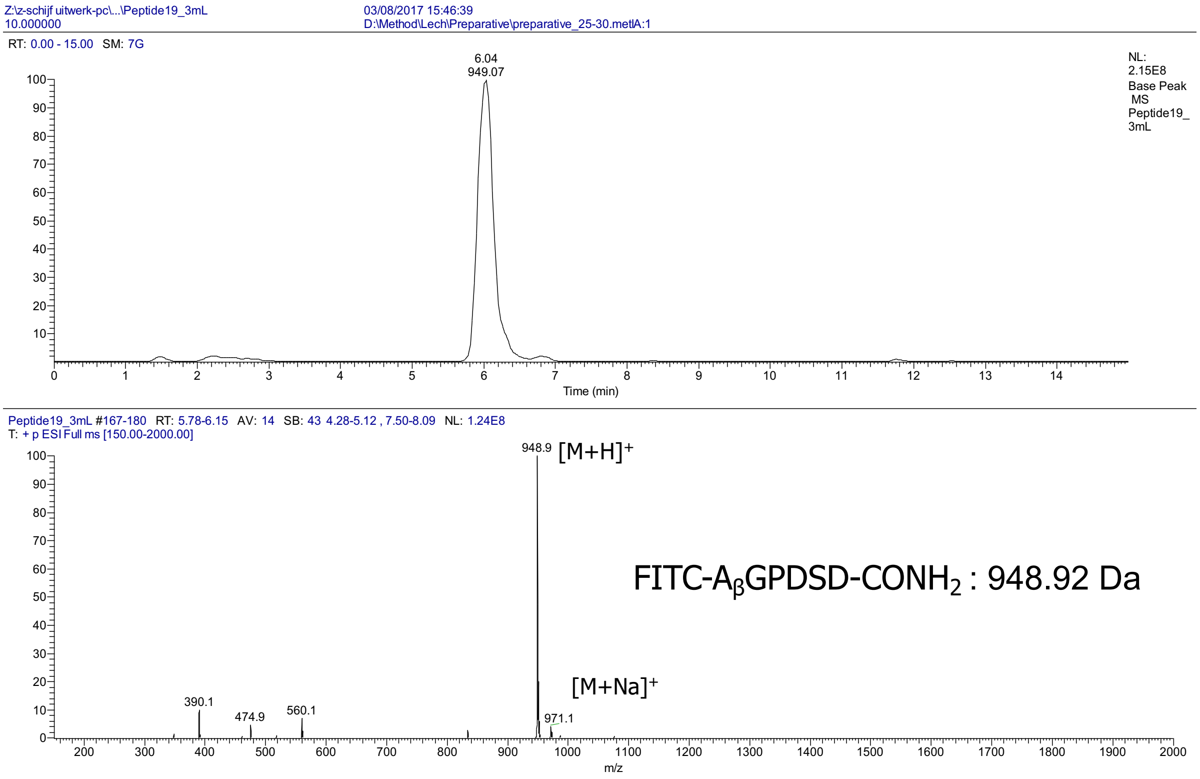


**Figure S13. Analytical LC/MS of the purified FITC-labelled peptide A$_\beta$GPDSD-CONH$_2$.** The graphs represent the base peak chromatogram (upper panel) and the mass spectrum (lower panel).

**HPLC Result**

| | | | |
|---|---|---|---|
| Product Name | Peptide 3 | | |
| Lot No. | P170802-GB599300 | | |
| Column | Column:250*4.6mm,Boston Green 0DS-AQ | | |
| Solvent A | 0.1% trifluoroacetic in 100% acetonitrile | | |
| Solvent B | 0.1% trifluoroacetic in 100% water | | |
| Gradient | | A | B |
| | 0.01min | 30% | 70% |
| | 25.00min | 55% | 45% |
| | 25.10min | 100% | 0% |
| | 30.00min | Stop | |
| Flow Rate | 1.0 ml/min | | |
| Wavelength: | 220nm | | |
| Volume: | 20μl | | |

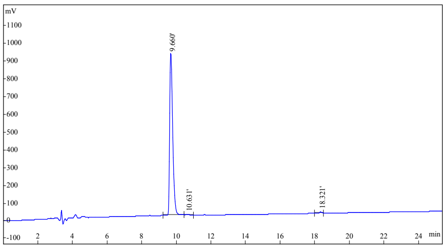

| Rank | Time | Conc | Area | Height |
|---|---|---|---|---|
| 1 | 9.660 | 98.8 | 11250354 | 922663 |
| 2 | 10.631 | 0.6965 | 79308 | 5700 |
| 3 | 18.321 | 0.4974 | 56632 | 6171 |
| Total | | 100.0000 | 11386294 | 934534 |

**Mass Spectrometry**

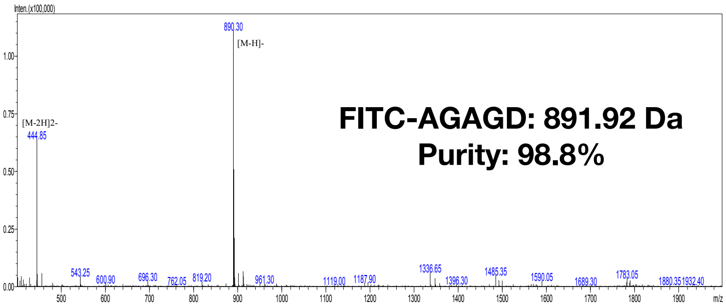


| Sample Description | | Instrument | SHIMADZU LCMS-2020 | | |
|---|---|---|---|---|---|
| Analyzed date: | 2017-8-8 | Probe: | ESI | Probe Bias: | +4.5kv |
| Analyst: | Wang | Nebulizer Gas Flow: | 1.5L/min | Detector: | 1.5kv |
| Sample: | Peptide 3 | CDL: | -20.0v | T. Flow: | 0.2ml/min |
| M.W.: | 891.92 | CDL Temp.: | 250 °C | B. Conc.: | 50%H2O/50%ACN |
| Lot. No.: | P170802-GB599300 | Block Temp.: | 200 °C | | |

**Figure S14. Analytical LC/MS of the purified peptide FITC-labelled AGAGD.** The graphs represent the HPLC chromatogram (left panel) and the mass spectrum (right panel). The Spectra were provided by GenicBio Limited.

**HPLC Result**

| | | | |
|---|---|---|---|
| Product Name | Peptide 17 | | |
| Lot No. | P171107-GB618641 | | |
| Column | 4.6×250mm, Kromasil 100-5 C18 | | |
| Solvent A | 0.1% trifluoroacetic in 100% acetonitrile | | |
| Solvent B | 0.1% trifluoroacetic in 100% water | | |
| Gradient | | A | B |
| | 0.01min | 30% | 70% |
| | 25.00min | 55% | 45% |
| | 25.10min | 100% | 0% |
| | 30.00min | Stop | |
| Flow Rate | 1.0 ml/min | | |
| Wavelength: | 220nm | | |
| Volume: | 5μl | | |

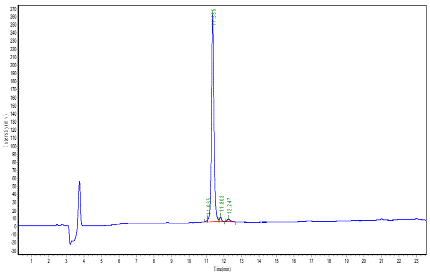

| Peak | Time | Height | Area | Conc. |
|---|---|---|---|---|
| 1 | 11.065 | 2440.145 | 10113.600 | 0.3871 |
| 2 | 11.355 | 256048.031 | 2513537.750 | 96.2167 |
| 3 | 11.803 | 5832.201 | 46426.023 | 1.7772 |
| 4 | 12.247 | 3151.185 | 42294.406 | 1.6190 |
| Total | | | | 100.0000 |

**Mass Spectrometry**

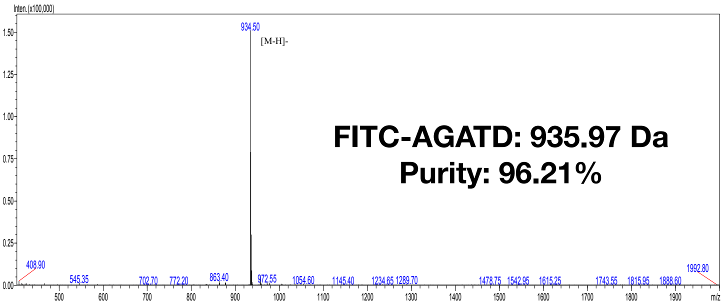


| Sample Description | | Instrument | SHIMADZU LCMS-2020 | | |
|---|---|---|---|---|---|
| Analyzed date: | 2017-11-9 | Probe: | ESI | Probe Bias: | +4.5kv |
| Analyst: | Wang | Nebulizer Gas Flow: | 1.5L/min | Detector: | 1.5kv |
| Sample: | Peptide 17 | CDL: | -20.0v | T. Flow: | 0.2ml/min |
| M.W.: | 935.97 | CDL Temp.: | 250 °C | B. Conc.: | 50%H2O/50%ACN |
| Lot. No.: | P171107-GB618641 | Block Temp.: | 200 °C | | |

**Figure S15. Analytical LC/MS of the purified peptide FITC-labelled AGATD.** The graphs represent the HPLC chromatogram (left panel) and the mass spectrum (right panel). The Spectra were provided by GenicBio Limited.

**HPLC Result**

| | | |
|---|---|---|
| Product Name | Peptide 3 | |
| Lot No. | P180814-GB672166 | |
| Column | Column:250*4.6mm,Boston Green 0DS-AQ | |
| Solvent A | 0.1% trifluoroacetic in 100% acetonitrile | |
| Solvent B | 0.1% trifluoroacetic in 100% water | |
| Gradient | | A | B |
| | 0.01min | 28% | 72% |
| | 25.00min | 53% | 47% |
| | 25.10min | 100% | 0% |
| | 30.00min | Stop | |
| Flow Rate | 1.0 ml/min | |
| Wavelength: | 220nm | |
| Volume: | 10μl | |

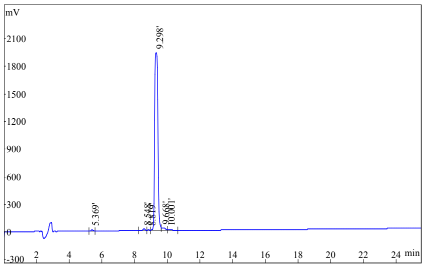


| Rank | Time | Conc | Area | Height |
|---|---|---|---|---|
| 1 | 5.369 | 0.4084 | 101693 | 13202 |
| 2 | 8.548 | 0.4372 | 108857 | 15802 |
| 3 | 8.819 | 0.1805 | 44935 | 5546 |
| 4 | 9.298 | 97.01 | 24158162 | 1928276 |
| 5 | 9.668 | 1.552 | 386357 | 22159 |
| 6 | 10.001 | 0.4083 | 101669 | 5818 |
| Total | | 100.0000 | 24901673 | 1990803 |

**Mass Spectrometry**

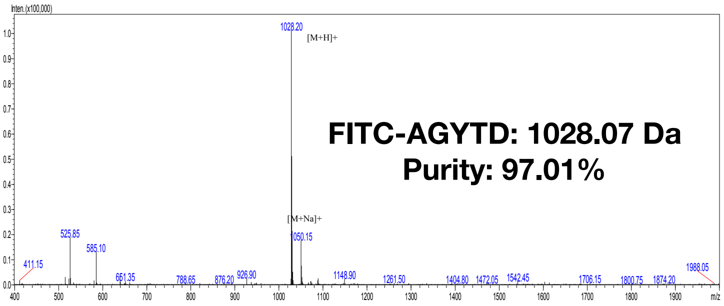


| | | | | | |
|---|---|---|---|---|---|
| Sample Description | | Instrument | SHIMADZU LCMS-2020 | | |
| Analyzed date: | 2018-8-29 | Probe: | ESI | Probe Bias: | +4.5kv |
| Analyst: | Wang | Nebulizer Gas Flow: | 1.5L/min | Detector: | 1.5kv |
| Sample: | Peptide 3 | CDL: | -20.0v | T. Flow: | 0.2ml/min |
| M.W.: | 1028.07 | CDL Temp.: | 250 °C | B. Conc.: | 50%H2O/50%ACN |
| Lot. No.: | P180814-GB672166 | Block Temp.: | 200 °C | | |

**Figure S16. Analytical LC/MS of the purified peptide FITC-labelled AGYTD.** The graphs represent the HPLC chromatogram (left panel) and the mass spectrum (right panel). The Spectra were provided by GenicBio Limited.

**HPLC Result**

| | | |
|---|---|---|
| Product Name | Peptide 16 | |
| Lot No. | P180814-GB672179 | |
| Column | 4.6×250mm, Cosmasil C18 Plus | |
| Solvent A | 0.1% trifluoroacetic in 100% acetonitrile | |
| Solvent B | 0.1% trifluoroacetic in 100% water | |
| Gradient | | A | B |
| | 0.01min | 27% | 73% |
| | 25.00min | 52% | 48% |
| | 25.10min | 100% | 0% |
| | 30.00min | Stop | |
| Flow Rate | 1.0 ml/min | |
| Wavelength: | 220nm | |
| Volume: | 10μl | |

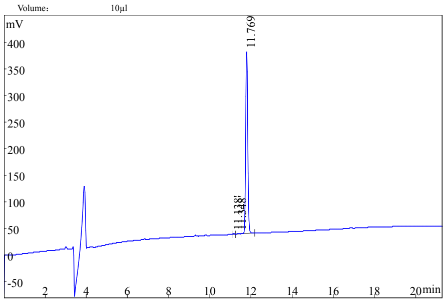


| Rank | Time | Conc. | Area | Height |
|---|---|---|---|---|
| 1 | 11.138 | 0.2751 | 5517 | 1092 |
| 2 | 11.348 | 0.2627 | 5268 | 794 |
| 3 | 11.769 | 99.46 | 1994706 | 341144 |
| Total | | 100.0000 | 2005491 | 343030 |

**Mass Spectrometry**

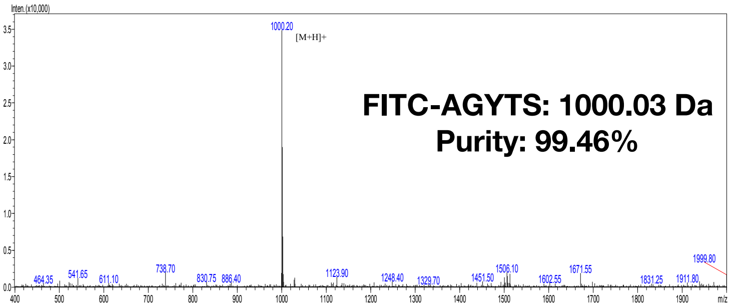


| | | | | | |
|---|---|---|---|---|---|
| Sample Description | | Instrument | SHIMADZU LCMS-2020 | | |
| Analyzed date: | 2018-8-20 | Probe: | ESI | Probe Bias: | +4.5kv |
| Analyst: | Wang | Nebulizer Gas Flow: | 1.5L/min | Detector: | 1.5kv |
| Sample: | Peptide 16 | CDL: | -20.0v | T. Flow: | 0.2ml/min |
| M.W.: | 1000.03 | CDL Temp.: | 250 °C | B. Conc.: | 50%H2O/50%ACN |
| Lot. No.: | P180816-GB672179 | Block Temp.: | 200 °C | | |

**Figure S17. Analytical LC/MS of the purified peptide FITC-labelled AGYTS.** The graphs represent the HPLC chromatogram (left panel) and the mass spectrum (right panel). The Spectra were provided by GenicBio Limited.

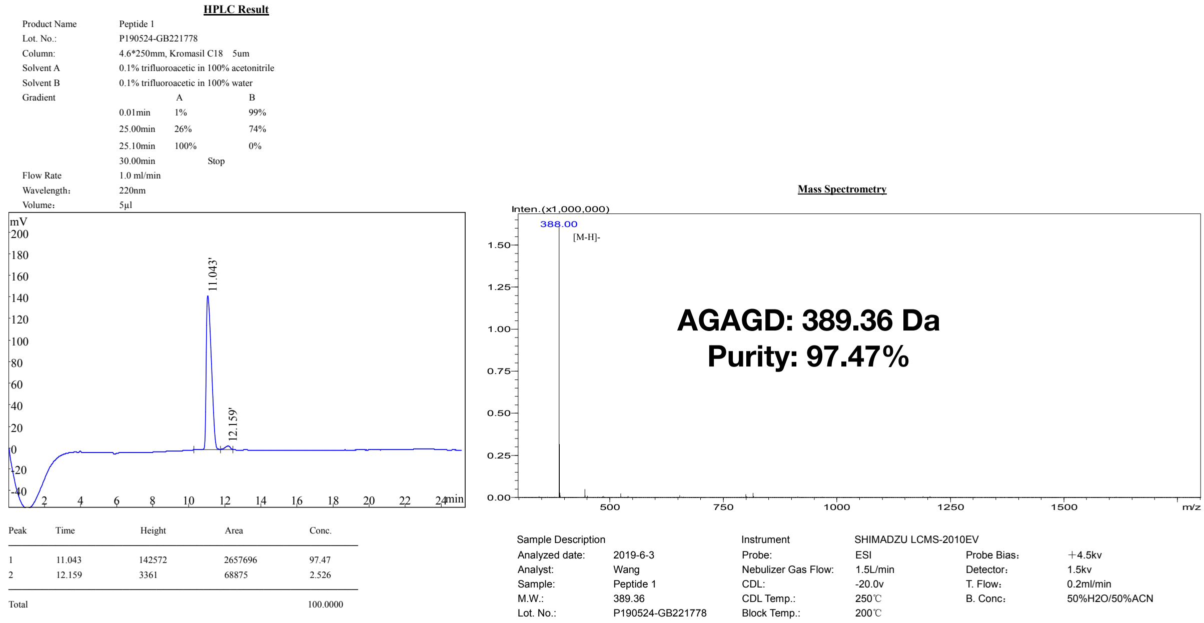


**Figure S18. Analytical LC/MS of the purified peptide AGAGD.** The graphs represent the HPLC chromatogram (left panel) and the mass spectrum (right panel). The Spectra were provided by GenicBio Limited.

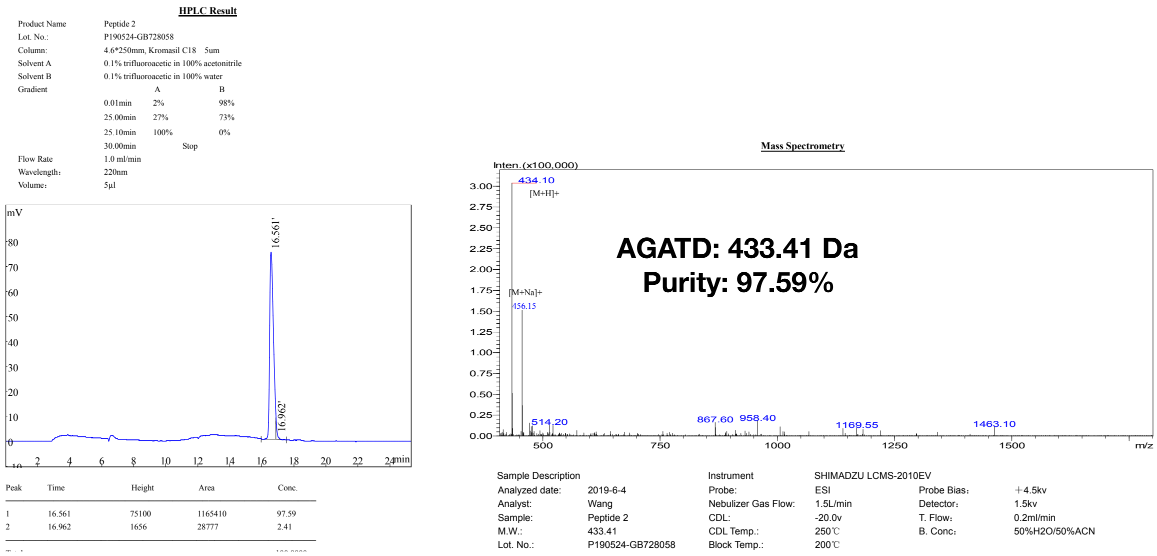


**Figure S19. Analytical LC/MS of the purified peptide AGATD.** The graphs represent the HPLC chromatogram (left panel) and the mass spectrum (right panel). The Spectra were provided by GenicBio Limited.

**HPLC Result**

| | | | |
|---|---|---|---|
| Product Name | Peptide 3 | | |
| Lot. No.: | P190524-GB728059 | | |
| Column: | 4.6*250mm, Kromasil C18 5um | | |
| Solvent A | 0.1% trifluoroacetic in 100% acetonitrile | | |
| Solvent B | 0.1% trifluoroacetic in 100% water | | |
| Gradient | | A | B |
| | 0.01min | 3% | 97% |
| | 25.00min | 28% | 72% |
| | 25.10min | 100% | 0% |
| | 30.00min | Stop | |
| Flow Rate | 1.0 ml/min | | |
| Wavelength: | 220nm | | |
| Volume: | 5μl | | |

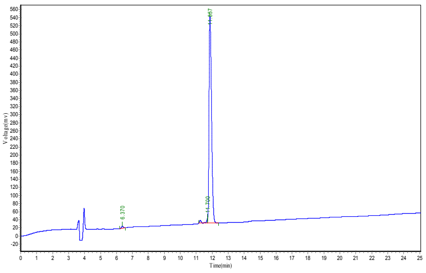


| Peak | Time | Height | Area | Conc. |
|---|---|---|---|---|
| 1 | 6.370 | 5391.330 | 42552.191 | 0.7383 |
| 2 | 11.700 | 11702.243 | 83957.383 | 1.4568 |
| 3 | 11.857 | 511295.594 | 5636764.500 | 97.8049 |
| Total | | | | 100.0000 |

**Mass Spectrometry**

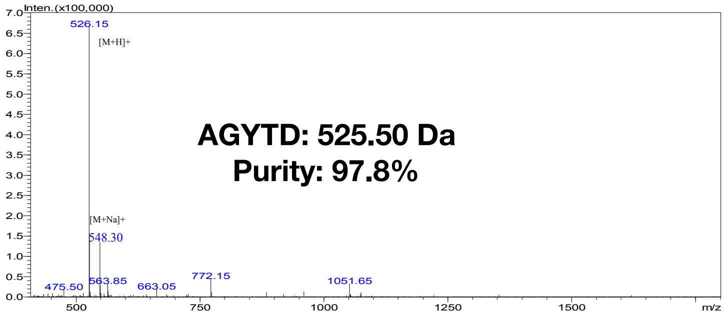


| Sample Description | | Instrument | SHIMADZU LCMS-2010EV | | |
|---|---|---|---|---|---|
| Analyzed date: | 2019-5-28 | Probe: | ESI | Probe Bias: | +4.5kv |
| Analyst: | Wang | Nebulizer Gas Flow: | 1.5L/min | Detector: | 1.5kv |
| Sample: | Peptide 3 | CDL: | -20.0v | T. Flow: | 0.2ml/min |
| M.W.: | 525.50 | CDL Temp.: | 250℃ | B. Conc: | 50%H2O/50%ACN |
| Lot. No.: | P190524-GB728059 | Block Temp.: | 200℃ | | |

**Figure S20. Analytical LC/MS of the purified peptide AGYTD.** The graphs represent the HPLC chromatogram (left panel) and the mass spectrum (right panel). The Spectra were provided by GenicBio Limited.

**HPLC Result**

| | | | |
|---|---|---|---|
| Product Name | Peptide 4 | | |
| Lot. No.: | P190524-GB728060 | | |
| Column: | 4.6*250mm, Kromasil C18 5um | | |
| Solvent A | 0.1% trifluoroacetic in 100% acetonitrile | | |
| Solvent B | 0.1% trifluoroacetic in 100% water | | |
| Gradient | | A | B |
| | 0.01min | 3% | 97% |
| | 25.00min | 28% | 72% |
| | 25.10min | 100% | 0% |
| | 30.00min | Stop | |
| Flow Rate | 1.0 ml/min | | |
| Wavelength: | 220nm | | |
| Volume: | 5μl | | |

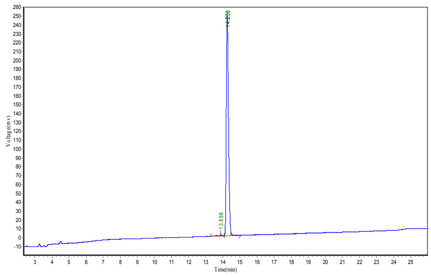


| Peak | Time | Height | Area | Conc. |
|---|---|---|---|---|
| 1 | 13.888 | 1490.446 | 18435.434 | 0.8194 |
| 2 | 14.255 | 248023.906 | 2216047.250 | 98.4958 |
| 3 | 14.255 | 3314.675 | 15406.315 | 0.6848 |
| Total | | | | 100.0000 |

**Mass Spectrometry**

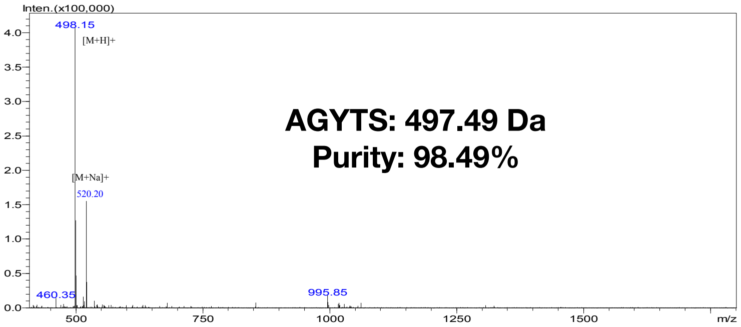


| Sample Description | | Instrument | SHIMADZU LCMS-2010EV | | |
|---|---|---|---|---|---|
| Analyzed date: | 2019-5-28 | Probe: | ESI | Probe Bias: | +4.5kv |
| Analyst: | Wang | Nebulizer Gas Flow: | 1.5L/min | Detector: | 1.5kv |
| Sample: | Peptide 4 | CDL: | -20.0v | T. Flow: | 0.2ml/min |
| M.W.: | 497.49 | CDL Temp.: | 250℃ | B. Conc: | 50%H2O/50%ACN |
| Lot. No.: | P190524-GB728060 | Block Temp.: | 200℃ | | |

**Figure S21. Analytical LC/MS of the purified peptide AGYTS.** The graphs represent the HPLC chromatogram (left panel) and the mass spectrum (right panel). The Spectra were provided by GenicBio Limited.

**Table S1. Statistics for the crystallographic data of *Mp*PBD in complex with four peptide ligands.**

| Data collection | FITC-A$_\beta$GPDSD | FITC-A$_\beta$DSTD | AGYTD | AGYTS |
|---|---|---|---|---|
| PDB accession code | 6X6Q | 6X6M | 6X5W | 6X5V |
| Space group | P1 | P1 | P1 | P1 |
| **Cell dimensions** | | | | |
| (a, b, c) (Å) | 50.2, 60.8, 63.7 | 50.1, 60.9, 63.8 | 51.1, 58.9, 63.2 | 49.8, 61,6, 63,4 |
| (α, β, γ) (°) | 66.6, 72.5, 68.6 | 66.4, 72.8, 68.3 | 62.5, 85.8, 65.9 | 119, 106, 93.3 |
| Resolution (Å) | 57.6 - 2 | 57.6 - 2 | 55.5-1.8 | 52.5 - 1.6 |
| No. of observations | 286662 | 287287 | 1100114 | 541809 |
| No. of unique | 41893 | 41931 | 53772 | 79169 |
| No. molecules/asymmetric unit | 1 | 1 | 1 | 1 |
| I/σI | 8.7 (4) | 9.4 (4) | 16.22 (3.6) | 12.10 (2.1) |
| $R_{merge}$ | 0.29 (0.63) | 0.2 (1.8) | 0.146 (0.809) | 0.069 (0.665) |
| CC(1/2) | 0.557 (0.21) | 0.61 (0.45) | 0.99 (0.87) | 0.99(0.76) |
| Completeness | 97.7 (91.5) | 98 (96.8) | 97.7 (94.4) | 96.9 (92.9) |
| Multiplicity | 6.8 (6.5) | 6.9 (6.8) | 20.5 (19.8) | 6.8 (6.9) |
| **Refinement** | | | | |
| Resolution (Å) | 57.6 - 2 | 57.6 - 2 | 47.3 - 1.8 | 52.5 - 1.8 |
| $R_{work}$/ $R_{free}$ (%) | 15.5/20.6 | 13.3/20.8 | 18.5/20.3 | 16.5/21.3 |
| No. of atoms | | | | |
| Amino acids/ions/water | 3792/16/655 | 3822/16/421 | 3838/22/544 | 3823/15/476 |
| B-factors (Å$^2$) | | | | |
| Amino acids/ions/water | 20.4/24.7/33.7 | 31.7/34.2/39 | 22.7/30.5/31.7 | 34.8/30.3/42.7 |
| **rms deviations** | | | | |
| **Bond lengths (Å)** | 0.021 | 0.017 | 0.01 | 0.002 |
| Bond angles (°) | 2 | 1.7 | 1.173 | 0.52 |
| Ramachandran statistics | | | | |
| Favored (%) | 98.4 | 98.2 | 97.2 | 98.02 |
| Outliers (%) | 0.99 | 0.79 | 0 | 0 |

**Table S2: Average $EC_{50}$ values calculated from the binding of FITC-labelled AGAGX and AGAXD peptides to *Mp*PBD determined by FP.** The strongest ligands from each of the two rounds of screening are bolded and underlined. Corresponding FP titration plots are shown in Figure S2 and S3.

| Peptides (AGAGX) | $EC_{50}$ (μM) | $EC_{50}$ 95% CI (μM) | Peptides (AGAXD) | $EC_{50}$ (μM) | $EC_{50}$ 95% CI (μM) |
|---|---|---|---|---|---|
| AGAGA | 28.4 | 2.8 to 286.4 | AGAAD | 0.47 | 0.38 to 0.59 |
| AGAGC | - | - | AGACD | 0.26 | 0.23 to 0.3 |
| **<u>AGAGD</u>** | **<u>3.2</u>** | **<u>2.1 to 5.0</u>** | AGADD | 1.1 | 0.67 to 1.9 |
| AGAGE | 38.9 | 4.3 to 349.8 | AGAED | 1.3 | 0.89 to 1.8 |
| AGAGF | 7.2 | 2.1 to 25.1 | AGAFD | 0.18 | 0.16 to 0.21 |
| AGAGG | - | - | AGAGD | 1.5 | 1.1 to 2.0 |
| AGAGH | - | - | AGAHD | 0.53 | 0.45 to 0.61 |
| AGAGI | 4.4 | 3.0 to 6.5 | AGAID | 0.64 | 0.52 to 0.77 |
| AGAGK | - | - | AGAKD | 1.3 | 1.0 to 1.6 |
| AGAGL | - | - | AGALD | 0.76 | 0.58 to 1.0 |
| AGAGM | - | - | AGAMD | 0.73 | 0.62 to 0.87 |
| AGAGN | 56.7 | 9.3 to 345 | AGAND | 0.56 | 0.50 to 0.62 |
| AGAGP | - | - | AGAPD | 4.6 | 2.5 to 8.6 |
| AGAGQ | - | - | AGAQD | 0.72 | 0.58 to 0.89 |
| AGAGR | - | - | AGARD | 1.2 | 1.0 to 1.3 |
| AGAGS | 18.5 | 5.9 to 57.7 | AGASD | 0.31 | 0.27 to 0.35 |
| AGAGT | 26 | 6.9 to 97.8 | **<u>AGATD</u>** | **<u>0.089</u>** | **<u>0.077 to 0.10</u>** |
| AGAGV | 507.9 | 4.6 to 55613 | AGAVD | 0.64 | 0.49 to 0.82 |
| AGAGW | 1302 | 0.0064 to 263960854 | AGAWD | 0.82 | 3.5e-005 to 18277 |
| AGAGY | 4.6 | 2.9 to 7.1 | AGAYD | 0.17 | 0.15 to 0.20 |

**Table S3: Average $EC_{50}$ values calculated from the binding of FITC-labelled AGXTD and AGYTX peptides to *Mp*PBD determined by FP.** The strongest ligands from each of these two rounds of screening are bolded and underlined. Corresponding FP titration plots are shown in Figure S4 and S5.

| Peptides (AGXTD) | $EC_{50}$ (µM) | $EC_{50}$ 95% CI (µM) | Peptides (AGYTX) | $EC_{50}$ (µM) | $EC_{50}$ 95% CI (µM) |
|---|---|---|---|---|---|
| AGATD | 0.07 | 0.059 to 0.083 | AGYTA | 0.056 | 0.044 to 0.071 |
| AGCTD | 0.023 | 0.017 to 0.031 | AGYTC | - | - |
| AGDTD | 0.076 | 0.057 to 0.099 | **<u>AGYTD</u>** | **<u>0.032</u>** | **<u>0.021 to 0.049</u>** |
| AGETD | 0.089 | 0.074 to 0.11 | AGYTE | 0.05 | 0.038 to 0.065 |
| AGFTD | 0.036 | 0.032 to 0.041 | AGYTF | 0.083 | 0.054 to 0.13 |
| AGGTD | 0.042 | 0.038 to 0.047 | AGYTG | 1.8 | 1.2 to 2.7 |
| AGHTD | 0.069 | 0.057 to 0.084 | AGYTH | 0.61 | 0.42 to 0.91 |
| AGITD | 0.06 | 0.054 to 0.067 | AGYTI | 0.062 | 0.047 to 0.081 |
| AGKTD | 0.081 | 0.066 to 0.099 | AGYTK | 6.8 | 3.5 to 13 |
| AGLTD | 0.071 | 0.060 to 0.084 | AGYTL | 0.58 | 0.35 to 0.96 |
| AGMTD | 0.053 | 0.046 to 0.061 | AGYTM | 0.25 | 0.15 to 0.42 |
| AGNTD | 0.061 | 0.054 to 0.068 | AGYTN | 0.26 | 0.14 to 0.33 |
| AGPTD | 0.092 | 0.079 to 0.10 | AGYTP | 34.2 | 0.15 to 7837 |
| AGQTD | 0.064 | 0.055 to 0.076 | AGYTQ | 0.075 | 0.061 to 0.092 |
| AGRTD | 0.072 | 0.060 to 0.086 | AGYTR | 1.6 | 1.1 to 2.2 |
| AGSTD | 0.073 | 0.064 to 0.081 | **<u>AGYTS</u>** | **<u>0.03</u>** | **<u>0.020 to 0.045</u>** |
| AGTTD | 0.063 | 0.057 to 0.070 | AGYTT | 0.12 | 0.077 to 0.20 |
| AGVTD | 0.064 | 0.057 to 0.071 | AGYTV | 0.12 | 0.080 to 0.17 |
| AGWTD | 0.023 | 0.019 to 0.029 | AGYTW | 18.8 | 0.023 to 15685 |
| **<u>AGYTD</u>** | **<u>0.029</u>** | **<u>0.026 to 0.031</u>** | AGYTY | 0.25 | 0.097 to 0.66 |

**Table S4. Ionic and hydrogen bonds at the protein-peptide interfaces. Length of the bonds are indicated.** Protein (left) and peptide (right) atoms involved in the interactions are listed.

| | *Mp*PBD | TPD | DSD | STD | YTD | YTS |
|---|---|---|---|---|---|---|
| **Ionic bonds** | | | | | | |
| | **Ca1** | Asp-O/2.7 Å | Asp-O/2.6 Å | Asp-O/2.7 Å | Asp-O/2.6 Å | Ser-O/2.6 Å |
| | **Ca1** | Asp-OXT/2.4 Å | Asp-OXT/2.5 Å | Asp-OXT/2.4 Å | Asp-OXT/2.6 Å | Ser-OXT/2.5 Å |
| | **Ca2** | Asp-OXT/2.5 Å | Asp-OXT/2.4 Å | Asp-OXT/2.5 Å | Asp-OXT/2.4 Å | Ser-OXT/2.4 Å |
| **Hydrogen Bonds** | | | | | | |
| **Peptide C terminus**<br>**1st position** | | | | | | |
| | **T331-O** | Asp-OXT/3.1 Å | Asp-OXT/2.9 Å | Asp-OXT/3.1 Å | Asp-OXT/2.8 Å | Ser-OXT/3 Å |
| | **V293-N** | Asp-O/3 Å | Asp-O/3 Å | Asp-O/2.8 Å | Asp-O/2.8 Å | Ser-O/3.1 Å |
| | **V293-O** | Asp-N/2.7 Å | Asp-N/2.6 Å | Asp-N/2.8 Å | Asp-N/2.6 Å | Ser-N/2.8 Å |
| | **G296-N** | - | Asp-OD2/3.5 Å | Asp-OD2/3.5 Å | Asp-OD2/2.8 Å | - |
| | **S295-OG** | - | Asp-OD1/3.2 Å | Asp-OD1/3.2 Å | Asp-OD1/2.5 Å | Ser-O/2.7 Å |
| **2nd position** | | | | | | |
| | **N333-N** | Pro-O/3.1 Å | Thr-O/2.9 Å | Thr-O/2.8 Å | Thr-O/3 Å | Thr-O/2.9 Å |
| | **N333-ND2** | - | Thr-OG1/2.8 Å | Thr-OG1/3.1 Å | Thr-OG1/2.8 Å | Thr-OG1/2.9 Å |
| **3rd position** | | | | | | |
| | **S295-N** | Thr-O/3 Å | Asp-O/3.3 Å | Ser-O/3.2 Å | Tyr-0/2.9 Å | Tyr-0/3 Å |
| | **S295-OG** | Thr-0/2.7 Å | - | - | - | - |
| | **S295-OG** | | - | - | Tyr-O/3 Å | Tyr-O/3.3 Å |
